# Investigating the composition of motor volleys directed to the hand through microneurography during transcranial magnetic stimulation of the motor cortex

**DOI:** 10.64898/2026.11.22.768460

**Authors:** Miriam Ceraman, Donggyu Rim, Brendan McCarthy, Gianni Sesa-Ashton, Leo Chen, Vaughan G Macefield, Andris Cerins

## Abstract

**Introduction:** Corticospinal volleys (CSV) induced by transcranial magnetic stimulation (TMS) of the primary motor cortex (M1) have been previously recorded via epidural spinal electrodes. Here we sought to determine whether individual components of CSVs – early waves evoked by direct excitation of pyramidal tract neurones (D-waves) and later waves generated by indirect excitation via interneurons (I-waves) – could be recorded from peripheral nerve in the upper arm via microneurography.

**Method:** A tungsten microelectrode was inserted into the ulnar or median nerve of the upper arm in eleven participants and directed into a fascicle supplying intrinsic hand muscles, confirmed by evoked muscle twitches during intraneural stimulation. TMS was applied to the contralateral M1 via a figure-of-eight coil at different intensities and current directions.

**Results:** CSV-like signals were successfully recorded in eight participants. Probable D-waves were elicited during stimulation with the coil in the latero-medial (LM) orientation, with probable early I-waves being evoked first during posterior-anterior (PA) directed current.

Anterior-posterior (AP) directed current produced variable I-wave recruitment. I-wave-like responses appeared in distinct clusters with median latencies of 1.6 ms, 3.0 ms, 4.5 ms, and 6.0 ms after the suspected D-wave elicited during higher-intensity LM stimulation, occurring at 11.2 ± 1.18 ms. These wave recruitment patterns were consistent with those of epidural recordings.

**Conclusion:** Intraneural microneurography can detect individual components of TMS-evoked CSVs, previously only possible via epidural electrodes. Recording D- and I-waves from peripheral nerves could lead to a deeper understanding of TMS neurophysiology, neuromodulation, and the neural substrates underlying MEP-based biomarkers.

**Highlights:**

- First recordings of TMS-evoked CSV components from peripheral nerve
- Volley components varied with current direction, matching epidural recording data
- Findings align with prior epidural electrode recordings of TMS-evoked volleys
- TMS CSV neurophysiology can now be studied in larger samples than before

## Introduction

Transcranial magnetic stimulation (TMS) of the motor cortex allows a non-invasive index of cortical activity that has substantially advanced our understanding of cortical neurophysiology [1] and has subsequently been applied to develop biomarkers for several neuropsychiatric conditions [2]. These advances have primarily been based on quantifying changes in the amplitude and latency of motor-evoked potentials (MEPs), which reflect the superimposed activity of multiple cortical and spinal circuits [3–6]. However, MEPs, whilst being a non-invasive method of recording, represent a compound cortical output made up of a series of distinct waves known as the corticospinal volley (CSV). These wave components may represent activity in dissociable neural populations [3, 7, 8] that are both differentially affected in biomarker assessment [4], and relevant to neuromodulation outcomes [9], but they cannot be resolved using the MEP. Recording the CSV directly might therefore provide richer data than the MEP and could improve interpretation of TMS probes of cortical activity. Invasive epidural recordings have revealed individual components of CSVs, but such data are rare and limited to patients with spinal epidural electrodes [10–13]. The limited participant pool and window of opportunity to record CSVs using epidural electrodes limits neurophysiological research, potentially hampering biomarker discovery.

When the primary motor cortex (M1) is stimulated with TMS at a sufficient, or ‘threshold’, intensity, a CSV descends through the (primarily) lateral corticospinal tract to activate lower spinal motoneurones [1]. The evoked action potentials then propagate along motor axons to produce MEPs, measurable via surface electromyography (sEMG) [5]. CSV composition depends on stimulus intensity and cortical current direction as determined by the orientation of a figure-of-eight TMS coil [3, 4, 6–8]. Higher-intensity stimulation with lateromedially (LM) directed current can produce a CSV that begins with the earliest volley component – a direct (D)-wave resulting from direct excitation of fast-conducting pyramidal tract neurones. The D-wave is usually followed by indirect (I)-waves arising from transsynaptic activation of pyramidal tract neurones [14–16]. Posterior-anterior (PA) stimulation at peri-threshold intensity tends to evoke early I-waves, or I1 and I2 waves at approximately 1.5 ms intervals, which begin around 1 to 2 ms after D-wave latency; these can be followed by later waves as stimulus intensity increases. Peri-threshold anterior-posterior (AP) stimulation tends to elicit later I-waves (I3 onward) commencing approximately 4 ms after D-wave latency. As AP stimulus intensity is increased, earlier and/or later waves can also be evoked [11, 13, 17].

TMS-evoked MEPs serve as real-time indicators of corticospinal output, with single-pulse (spTMS) and paired-pulse TMS (ppTMS) paradigms assessing cortical excitability, intracortical facilitation and inhibition [5, 18]. This has led to the development of potential TMS biomarkers for several neurological and psychiatric conditions [2, 19]. Indexing the underlying CSV profiles of these TMS measures in neuropsychiatric conditions may offer further insight into the underlying neural mechanisms and improve the accuracy of these potential biomarkers. But there are several limitations in recording them. As mentioned above, invasive epidural recordings have restricted cohorts; sEMG (though non-invasive), is not suited to isolating individual motor unit (MU) activity [3, 5] and is a proxy measure of the earliest component of a CSV; finally, Intramuscular EMG (IM-EMG) is time-consuming [20–22] as it requires 100-200 TMS sweeps to populate the histogram and obtain a representation of the CSV, and is prone to interference from overlapping MU potentials [23] that cannot be corrected even with discriminator software.

Microneurography [24–26] may partly address these limitations. The procedure involves inserting a fine tungsten microelectrode percutaneously into a fascicle of a peripheral (or cranial) nerve, which allows one to directly record action potentials generated by single sensory or motor axons, or groups of axons, in a minimally invasive and safe fashion. However, only one study (to the best of the authors’ knowledge) has used this approach to record TMS-evoked motor volleys: McNulty et al. (2002) [22] recorded motor volleys to the intrinsic hand muscles via a microelectrode inserted into the ulnar nerve in the upper arm before and during progressive ischaemic block of the hand. Both motor volleys and sensory volleys associated with evoked muscle twitches could clearly be recorded. However, they did not identify individual CSV components nor was there preferential activation of D- and/or I- waves through changes in cortical current direction. Nevertheless, this study confirmed that motor volley signals can be detected in peripheral nerves using microneurography.

We reasoned that identifying individual components of the CSV with microneurography might be possible and could provide rich information on TMS neural recruitment without requiring epidural electrode recordings. Therefore, the current study aimed to use microneurography, combined with changes in TMS current direction, to identify individual D- and I-waves comprising multi-unit motor volleys. We hypothesised that we would be able to record volley-like features from the peripheral nerve via microneurography. We further hypothesised that recruitment of D- and I-waves would depend on current direction and stimulation intensity as reported in previous studies involving epidural electrode recordings.

## Methods

Healthy participants were recruited, with standard TMS safety screening applied during the process [27]. The study was approved by the Monash University Human Research Ethics Committee (MUHREC 45198) and conformed to the Declaration of Helsinki. Informed, written consent was obtained from all participants.

### Transcranial Magnetic Stimulation

Motor cortex hotspots (where the largest single-trial MEPs were elicited) were located using standard search procedures [28] and recorded in a neuronavigational system brain template (TMS Navigator, Localite, Germany). A patient tracker on the forehead and a coil tracker ensured consistent coil positioning. PA stimulation hotspots were used for LM and AP stimulation. TMS pulses were delivered via a figure-of-eight MagVenture Cool B65 TMS coil (65 mm wing diameter, MagVenture, Denmark), with intensity expressed as a percentage of maximum stimulator output (%MSO; X100 standard mode) or %MSO + 40% (X100 power mode). Blocks of 5-10 pulses were delivered using AP, PA, or LM directed current with 4-8 s between pulses, during slight voluntary contraction, with rest between blocks to avoid fatigue.

Standard surface Ag/AgCl electrodes (Ambu, Ballerup, Denmark) were placed over the first dorsal interosseous, abductor digiti minimi, and abductor pollicis brevis. Electromyographic activity was amplified and filtered (2,000×, bandpass 0.01-5 kHz), sampled at 20 kHz, and acquired via a computer-based data acquisition and analysis system (PowerLab 8/35 hardware and LabChart 8 software (ADInstruments, Sydney, Australia)).

### Microneurography

The right ulnar or median nerve in the medial bicipital groove was located using a skin surface probe delivering brief cathodal pulses (0.2 ms, 3-10 mA, 1 Hz) from an isolated stimulator (Stimulus Isolator, ADInstruments, Sydney) to the medial upper arm, with a surface electrode placed opposite on the lateral upper arm serving as the anode. The site at which the strongest twitches of intrinsic hand muscles were evoked at the lowest current was marked on the skin. A sterile insulated tungsten microelectrode (FHC, USA) was then inserted through the alcohol-swabbed skin at the marked site, and an uninsulated tungsten microelectrode inserted subdermally 1-2 cm away from the recording microelectrode acted as the reference electrode. By recording from the upper rather than the lower arm, TMS-evoked volleys would theoretically have less time to disperse, thereby preserving individual volley components.

The recording electrode tip was manually manipulated toward the nerve, guided by response to low-intensity electrical stimuli (0.01 to 1 mA, 0.2 ms, 1 Hz) delivered through the microelectrode. Muscle twitches evoked at ≤ 0.02 mA indicated that the electrode tip had entered the nerve close to motor axons supplying hand muscle [22]. Evoked muscle spindle afferent activity from tendon percussion or stretching the target muscle was also used as an indication that the electrode tip had reached the nerve [29].

Once the electrode tip had been placed in the nerve, TMS was delivered while time-locked recordings were obtained from the nerve (via microneurography) and from the skin (via surface electrode) over the hand muscle.

Neural activity was amplified and filtered (20,000 x, bandpass 0.3-5 kHz; NeuroAmpEX, ADInstruments, Sydney, Australia), sampled at 20 kHz, and acquired via a computer-based data acquisition and analysis system (PowerLab 8/35 hardware and LabChart 8 software (ADInstruments, Sydney, Australia).

### CSV recordings and Data Analysis

Nerve and EMG signals were exported to MATLAB (R2023b) for baseline correction and averaging. Raw and acausally filtered (second-order Butterworth bandpass 0.3–3 kHz) nerve signals were visualised from −10 to +20 ms relative to the TMS pulse, with the EMG response being displayed from −10 to +50 ms following the TMS pulse. The number of sweeps per averaged plot ranged from 2 to 66, with a median of 10 sweeps. These plots were used to manually identify any peaks and peak latencies of volley-like features observed in the nerve signal (Figure 1). We subtracted latencies of any potential LM-stimulated D-waves from the latencies of waves evoked by PA and AP current direction, to ascertain the latency differences of recruited I-waves relative to the D-wave.

**Figure 1.**
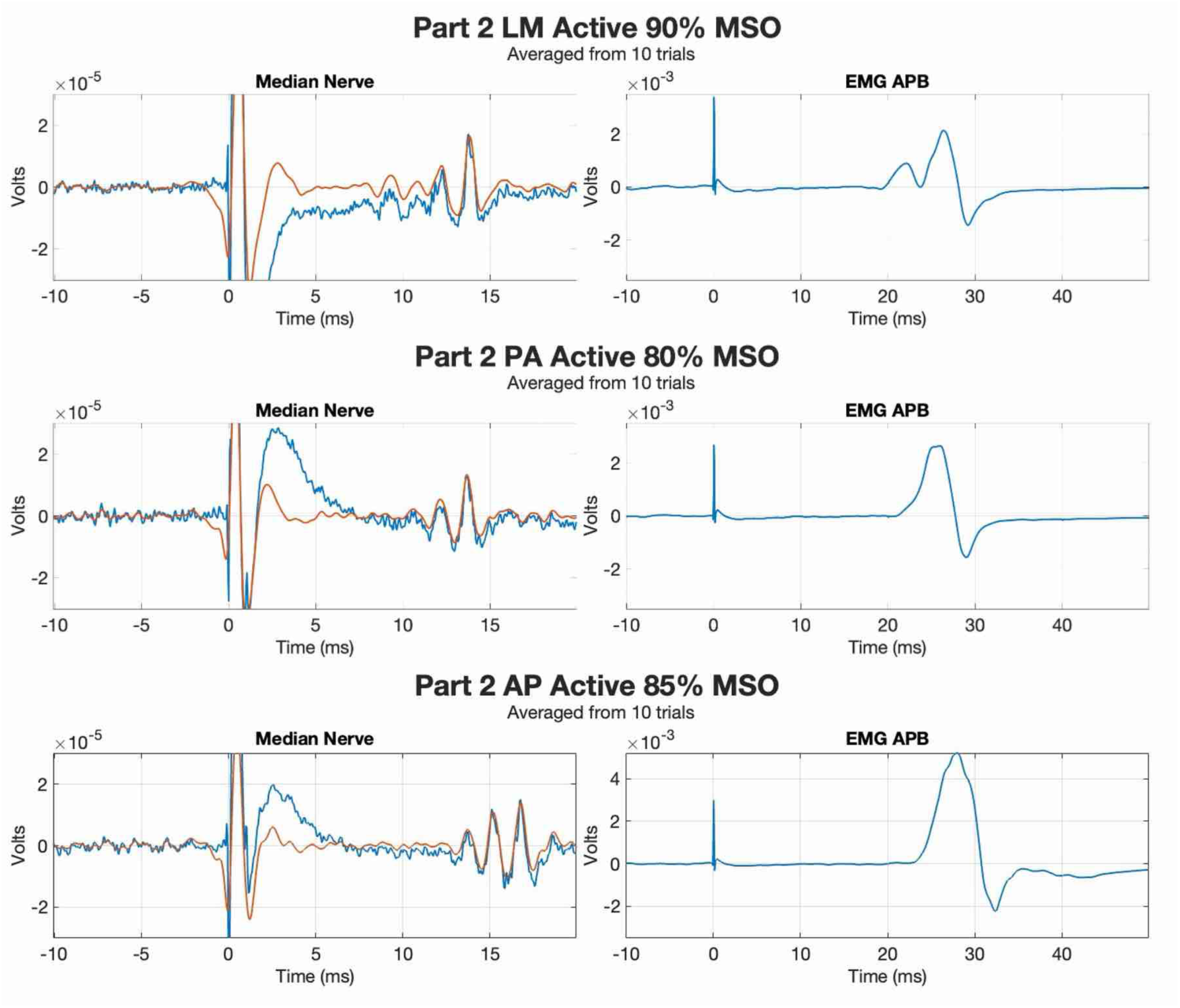
TMS-evoked volleys and their D- and I-waves recorded during LM, PA, and AP current directions in Participant 2. TMS-evoked volleys in the nerve signal (left column) and their respective motor evoked potentials in the surface EMG signal (right column), grouped by TMS cortical current direction, recorded in Participant 2. TMS pulses were delivered at Time Zero (as seen in both the Median Nerve and EMG APB recording at 0 ms). Part = Participant, Trial = a single TMS pulse; PA = Posterior-Anterior; LM = Latero-Medial; AP =Anterior-Posterior; APB = abductor pollicis brevis. Note the different time axis for the nerve and EMG traces.

## Results

Seventeen participants were recruited. In eleven participants (4 male, 7 female) aged 22-27 (24.8±1.9, mean±SD years), we were able to insert a microelectrode into a fascicle of the median or ulnar nerve in the upper arm and make recordings during TMS. Eight of these recordings (henceforth numbered participants 1, 2, 3, 4, 5, 6, 7, and 8) appeared to contain TMS-evoked motor volleys, while three did not.

### Timing of Features Observed in Nerve Signals

We observed a series of deflections in the nerve signal that occurred at approximately 1 to 2 ms intervals, consistent with D and I wave intervals reported in epidural recordings (Figure 1). Across all current directions and stimulus intensities, on average the first volley-like features in the nerve signals appeared 12.73 ms (SD = 1.35) after the TMS pulse while average MEP latency was 21.25 ms (SD = 2.17). We present visualisations of the nerve and EMG signals for all eight participants in the supplementary material.

In each participant, the highest stimulation intensity used during LM current direction produced the shortest latency responses in that particular individual, comprising an average of three volley components separated by approximately 1.5 ms. Interestingly, for participant 2, one component (a probable I1 wave) was present during LM stimulation at 90% but unexpectedly disappeared when stimulus intensity was increased to 95%, (Supplementary: Participant 2, Current Direction LM, MSO% 90-95%).

In four participants (1, 3, 4, 8), the latencies of the first deflection in the nerve trace were similar for PA and LM stimulation, potentially indicating that TMS had not elicited a D- wave. The earliest peak latencies for PA and LM stimulation were separated by at least 1 ms in the remaining four participants (2, 5, 6, 7), suggesting that TMS had possibly elicited a D- wave. For these participants, we subtracted the potential D-wave latencies from the latencies of responses to PA and AP stimulation to account for individual differences in the length of the neural pathway from cortex to our nerve recording site. When looking at all responses in these data, the peaks in the nerve recordings appeared in five clusters (Figure 2). Median latencies of these clusters were 0.2 ms, 1.6 ms, 3.0 ms, 4.5 ms, and 6.0 ms.

**Figure 2.**
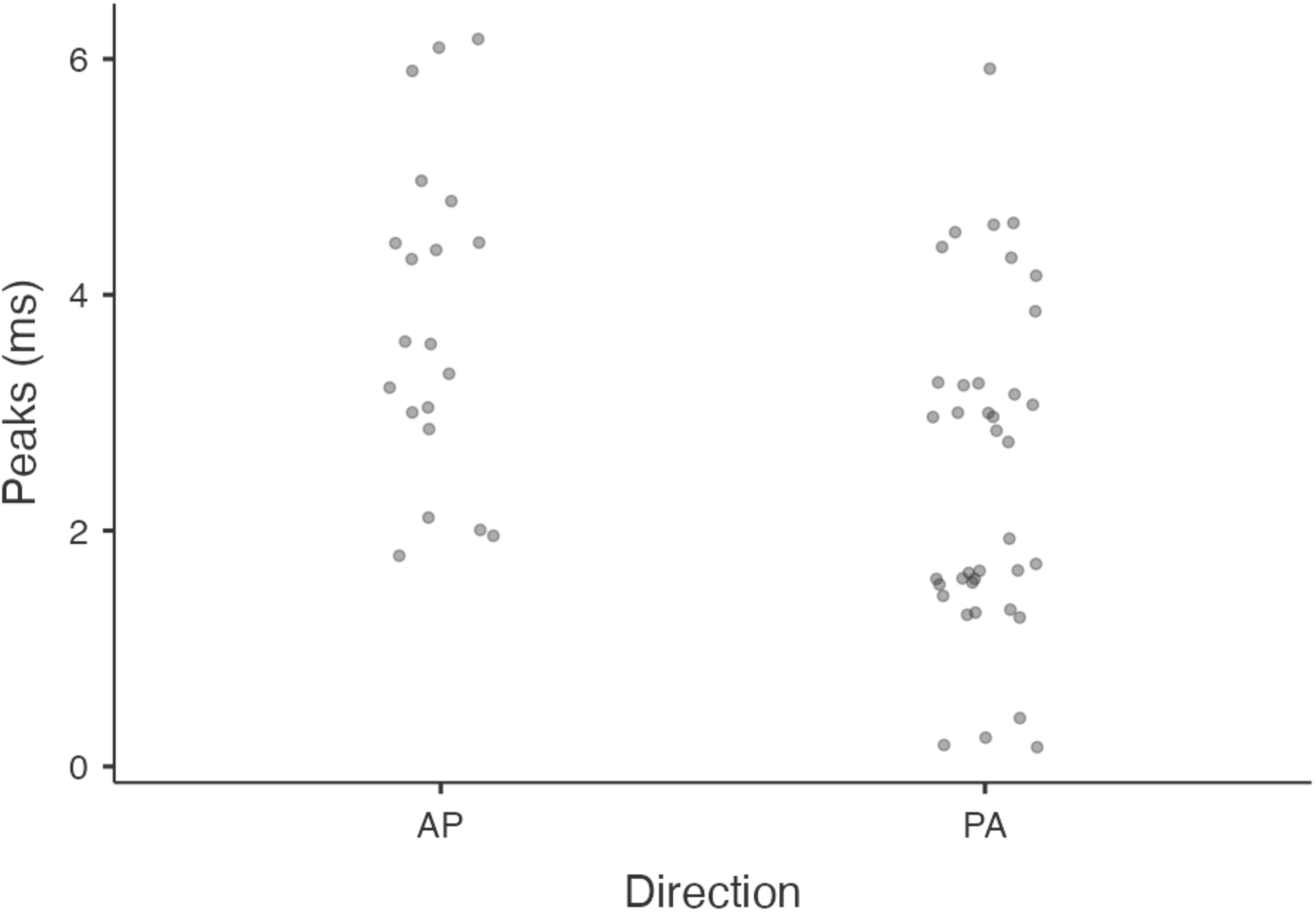
Timing of observed peaks in nerve signals. Points represent timing of observed peaks in each participant’s averaged nerve signals with the earliest peak detected during high intensity lateromedially directed stimulation subtracted. Data come from the four participants whose earliest LM latency was more than 1 ms earlier than their lowest intensity PA latency. Note clustering at approximate indirect wave intervals. PA = Posterior-Anterior; AP = Anterior-Posterior.

### PA stimulation

We found volley-like features in the nerve signal in eight participants during PA stimulation. In those with D-wave-like responses during LM stimulation, PA stimulation at the lowest intensity that produced volley-like features in each individual elicited peaks approximately 1.7 ms after their shortest LM latency, with smaller numbers of peaks occurring 2 to 4 ms later. At higher stimulus intensities, D-wave-like responses were occasionally evoked, and one 6 ms peak was observed (see Figure 2).

During PA stimulation at lower stimulus intensities, two participants (6 and 7) recruited a suspected I1 wave first, with latency approximately 1-1.5 ms longer than that of the earliest wave recruited during LM. Participant 2 recruited possible I1, I2, and I3 waves at the same time during 80%MSO. Participant 5 recruited a potential I2 wave, which appeared 2-3 ms after the earliest component they recruited during LM stimulation.

Four participants (1, 3, 4 and 8) saw the earliest components recruited during PA occurring at similar latencies to the waves recruited during their LM stimulation. It is possible that we did not recruit a D-wave for these participants during LM stimulation, and that the earliest component we are seeing in both their LM and PA stimulation is potentially an I1 wave.

As stimulus intensity increased, Participant 6 showed the recruitment of an earlier, potentially D-wave, followed by an I2 wave. Participant 7 showed the recruitment of suspected I2 waves and onwards as stimulus intensity increased. Participant 5 recruited an I1 wave after their I2 wave, then an I3 wave.

### AP stimulation

We found volley-like features in the nerve signal in six participants (2, 4, 5, 6, 7, 8) during AP stimulation. In those with D-wave-like responses during LM stimulation, most peaks occurred approximately 3.3 and 4.5 ms later, with other peaks approximately 2 and 6 ms later (Figure 2).

During AP stimulation at lower stimulus intensities, three participants (6, 7 and 8) recruited suspected early I-waves first, with Participant 7’s first recruited wave at 1-1.5 ms after the earliest component recruited during LM stimulation, and Participants 6 and 8 recruiting a wave where I2 waves would be expected. As stimulus intensity increased, Participants 6 and 8 showed an I3 wave recruited, whilst Participant 7 showed the recruitment of an I2 wave.

Participant 8’s results are based on the likelihood that the earliest component observed during LM stimulation, which reflected results obtained during PA stimulation, was not a D-wave, but an I1 wave.

Two participants (2 and 5) recruited multiple I-waves at a time during lower stimulus intensity. Participant 2 recruited a suspected I3 and I4 wave first, and as stimulus intensity increased, recruited a probable I5 wave. Participant 5 recruited I2, I3, and I4 waves at the same time during 90%MSO, and this stayed consistent as stimulus intensity increased.

Participant 4 recruited a single I3 wave first. As the stimulus increased, I2 and I4 waves were recruited simultaneously. This assumes that the earliest recorded component during LM stimulation, which mirrored results obtained during lower intensity PA stimulation, was an I1 wave and not a D-wave.

## Discussion

To the best of our knowledge, this is the first report to show that microneurography can be used to detect individual components of the CSV, i.e., D- and I-waves from the peripheral nerves in healthy people. Our results agree with our hypotheses and demonstrate that distinct CSV components can be identified using microneurography. We observed TMS-evoked volley-like signals occurring at 1-2 ms intervals. The latency of the first evoked component of the volley was approximately 60% of MEP latency, consistent with expected timing of signals travelling from the motor cortex to our recording site in the upper arm and onward to the hand. Our recordings also displayed intensity- and direction-dependent variation similar to previous work using epidural electrodes [8, 10–13]. Specifically, with reference to LM- determined estimates of D-wave latency, PA stimulation predominantly recruited earlier I- waves as the first component, aligning with known recruitment patterns in epidural recordings. As per the literature, I-wave recruitment was more variable during AP stimulation [7, 11, 17]. All eight participants’ PA recordings were consistent with epidural electrode findings, which show initial recruitment of earlier I-waves. Among the six participants that showed volleys during AP stimulation, three recruited early I-waves first, two recruited multiple waves at the same intensity, and one participant recruited an I3 wave first.

The ability to record directly from a peripheral nerve brings several advantages over previously available methods. To date, epidural recordings have provided the most direct information on the TMS-evoked CSV, but reliance on participants with implanted epidural electrodes has generally restricted study sample sizes to 1-10 participants [30]. Our observation of volley features in a peripheral nerve via microneurography makes it possible to expand the participant pool, making further investigation in larger numbers of participants more practically achievable. Additionally, unlike IM-EMG, which requires 100-200 sweeps to record the components of a CSV, our microneurography recordings used a median of 10 sweeps per condition.

Recording D- and I-waves from peripheral nerves could lead to a deeper understanding of the neural substrates underlying existing MEP-based biomarkers. One example of a biomarker is short-interval cortical inhibition (SICI) involving a subthreshold conditioning pulse followed by a suprathreshold test pulse at an interstimulus interval of 1-6ms, resulting in an inhibited MEP [31]. It is a measure of GABA-A mediated inhibition [31, 32]. One example of how SICI is used as a biomarker include that unmedicated patients with Parkinson’s disease show a reduction in SICI [33], indicating reduced motor cortex inhibition; however, when patients began taking medication, they showed increased SICI correlating with improved motor skills [33]. Conversely, it has been found that people with treatment-resistant major depressive disorder (MDD) had higher SICI compared to non-treatment-resistant MDD and healthy controls [34–37]. Importantly for the present work, epidural recordings suggest SICI exclusively inhibits the later components of the motor volley [38, 39] which means inter- individual differences in the TMS-evoked CSV influence the level of assessed inhibition [40, 41]. Thus, our proposed method of microneurography to directly record the TMS-evoked CSV and its D- and I-waves may provide a more accurate assessment of cortical inhibition via SICI, and potentially provide more detailed information for other TMS biomarkers. SICI is one of several existing TMS biomarkers, so there may be a wide range of potential applications for this research.

Access to previously underexplored details of the CSV might shed light on the neural correlates of reduced or amplified inhibition seen in several neurological and psychiatric conditions, and potentially increase understanding of underlying pathophysiological mechanisms that might be therapeutic targets. Further, given that later I wave recruitment is associated with TBS-induced neuromodulation outcomes, and with learning [9, 42–45], being able to record the CSV may help researchers better understand the organisation of cortical microcircuits recruited and modulated by TMS. Overall, recording D- and I-waves from peripheral nerves could lead to a deeper understanding of TMS neurophysiology, neuromodulation, and the neural substrates underlying existing MEP-based biomarkers.

### Limitations

There are several limitations to our work that must be acknowledged. Firstly, although the properties of our suspected D-waves were largely consistent with known direction- and intensity-dependent effects on CSV recruitment, we did not apply transcranial electrical stimulation [6] and cannot be absolutely certain that there were D-waves in our signal. Also, we did not appear to evoke D waves in four participants, possibly as a result of insufficient stimulus intensity.

Secondly, we did not observe volley-like features in the nerve signals in three participants, potentially due to the microelectrode tip not being close enough to the motor axons supplying intrinsic muscles of the hand. This would mean that evoked CSVs could not be recorded, though we were able to confirm afferent neural signals from muscle stretch or tendon percussion. In one participant, a volley component (a likely I1 wave in participant 2) unexpectedly disappeared as LM stimulation intensity was increased from 90 to 95 %MSO. This observation is difficult to explain. If specific groups of axons carry I1 waves it is possible the electrode tip may have moved away from these axons, potentially causing the I1 wave to disappear in the recorded signal. In addition, we frequently observed MEPs in hand muscles without detecting corresponding CSV components in the nerve signal, despite evidence (from muscle stretch and tendon percussion [46]) that the electrode tip had entered the nerve. It is possible that the high impedance of the microelectrodes, sometimes used to achieve remarkable single axon discrimination in microneurography, may not have detected neural traffic in axons further from the recording tip. In future research, lower-impedance electrodes might be able to detect signals from a larger volume of motor axons within a fascicle.

Thirdly, we did not apply motor thresholding but rather used raw stimulator output to express stimulation intensity and cannot interpret our data with the cross-participant comparison that percent motor threshold provides. Future research should establish motor threshold to allow greater comparability of stimulation intensity across participants.

Finally, in seeking to increase stimulus intensity, we occasionally used the stimulator’s power mode which also increases pulse width. There is some evidence that greater pulse width can affect neural recruitment, particularly with AP current [47], and acknowledge the possibility that greater pulse width in power mode may have produced greater TMS recruitment of cortical neurons.

## Conclusion

For the first time, we detected TMS-evoked volley-like signals in the peripheral nerve utilising microneurography. The features occurred at 1-2 ms intervals and displayed intensity- and direction-dependent variation similar to previous work using epidural electrodes. These results provide support for microneurography’s potential as a tool by which D- and I-waves can be recorded. This could pave the way for future studies, improving our understanding of the neurophysiology of the motor system and TMS markers in neuropsychiatric and neurological conditions.

