## supplementary material for "Investigating the composition of motor volleys directed to the hand through microneurography during transcranial magnetic stimulation of the motor cortex"

**Visualisation of the microneurography set up for CSV recording**

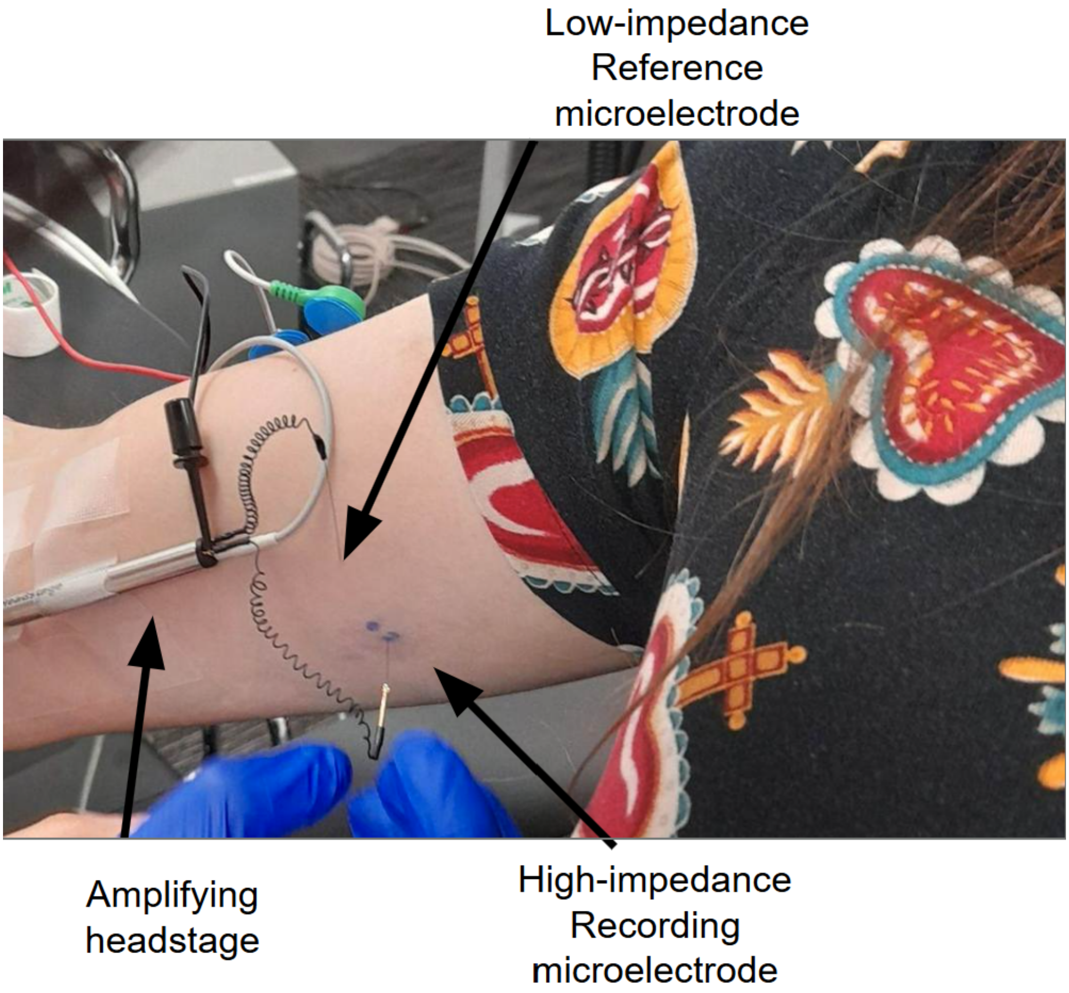


*Supplementary Figure 1. The microneurography set-up to record TMS-evoked volleys and their D- and I-waves during LM, PA, and AP current directions.* The amplifying headstage amplifies the signal being recorded at the site by the high-impedance recording microelectrode. A low-impedance reference electrode is placed a few centimetres away to serve as a stable ground or electrical baseline.

**Visualisation of the CSV recordings organised by participant and current direction**

The following supplementary figures contain visualisations of all volley-like activity recruited in all participants, ordered by participant number. Within each set of an individual participant's volley recordings, the images are ordered by current direction (LM, PA, AP). Then, each current direction is ordered by increasing %MSO. This material can be used in conjunction with the results section of this paper to better understand the distribution patterns of D- and I-waves per TMS current direction per individual participant.

TMS-evoked volleys in the nerve signal (left column) and their respective motor evoked potentials in the surface EMG signal (right column), grouped by TMS cortical current direction. Plots from all participants at stimulation intensities where volley-like features were visible. Part = Participant; PA = Posterior-Anterior; LM = Latero-Medial; AP =Anterior-Posterior; APB = abductor pollicis brevis. Note the different time axis for the nerve and EMG traces. Stimulation intensities expressed as % maximal stimulator output for MagPro X100 standard mode, .1 suffix indicates use of power mode.

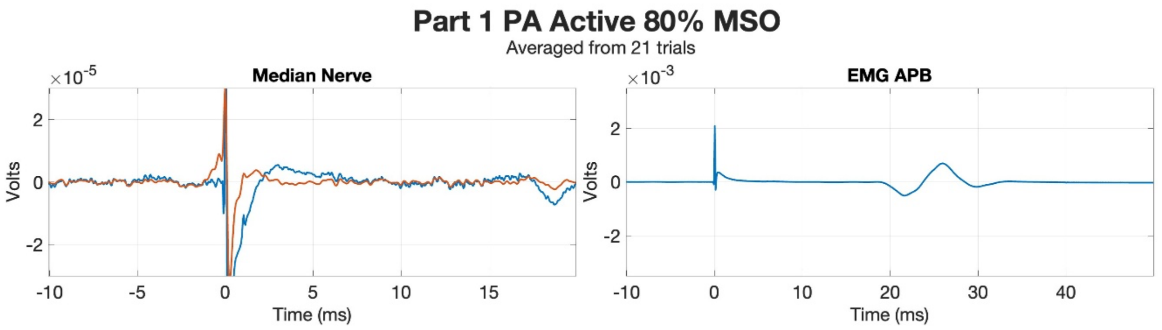
Part 1 PA Active 80% MSO
Averaged from 21 trials
Median Nerve
EMG APB
Volts
Time (ms)

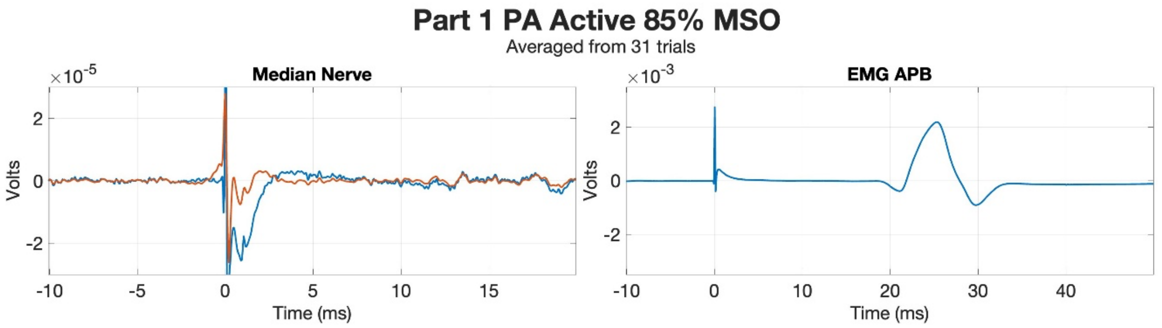
Part 1 PA Active 85% MSO
Averaged from 31 trials
Median Nerve
EMG APB
Volts
Time (ms)

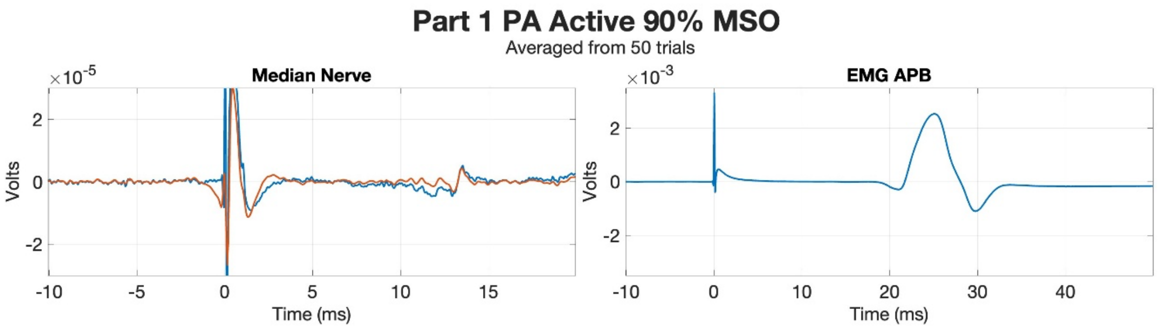
Part 1 PA Active 90% MSO
Averaged from 50 trials
Median Nerve
EMG APB
Volts
Time (ms)

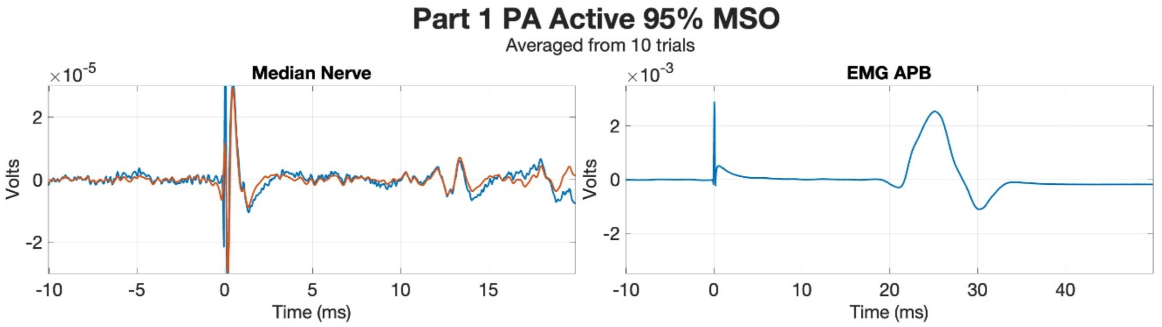
Part 1 PA Active 95% MSO
Averaged from 10 trials
Median Nerve
EMG APB
Volts
Time (ms)

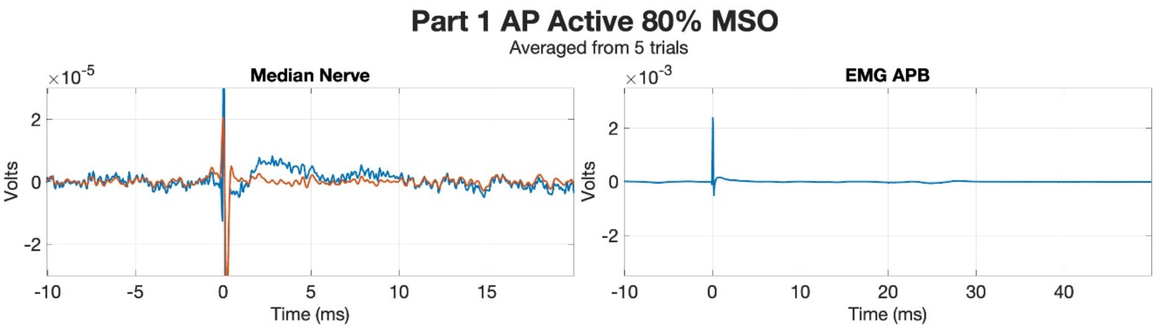
Part 1 AP Active 80% MSO
Averaged from 5 trials
Median Nerve
EMG APB
Volts
Time (ms)

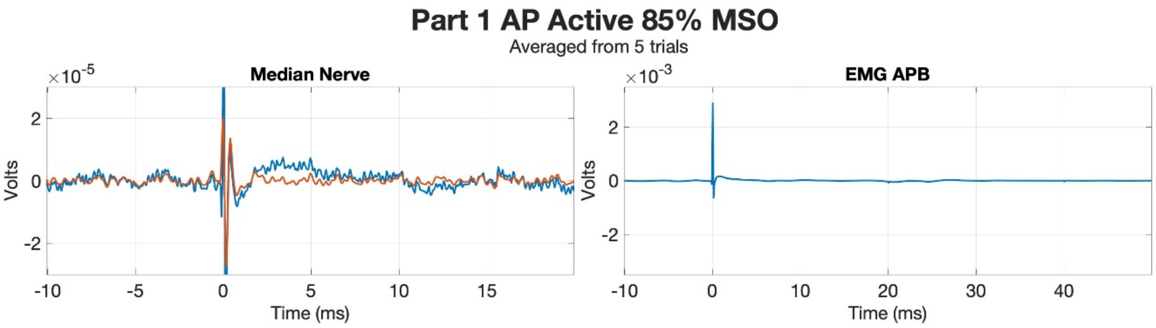
Part 1 AP Active 85% MSO
Averaged from 5 trials
Median Nerve
EMG APB
Volts
Time (ms)

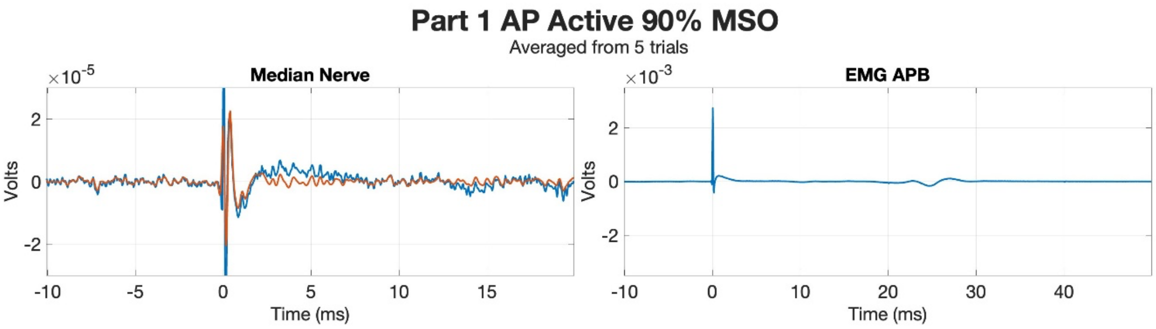
Part 1 AP Active 90% MSO
Averaged from 5 trials
Median Nerve
EMG APB
Volts
Time (ms)

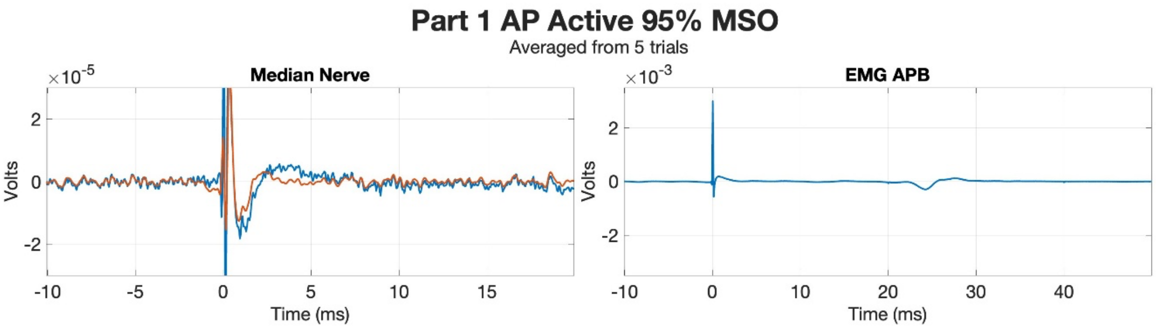
Part 1 AP Active 95% MSO
Averaged from 5 trials
Median Nerve
EMG APB
Volts
Time (ms)

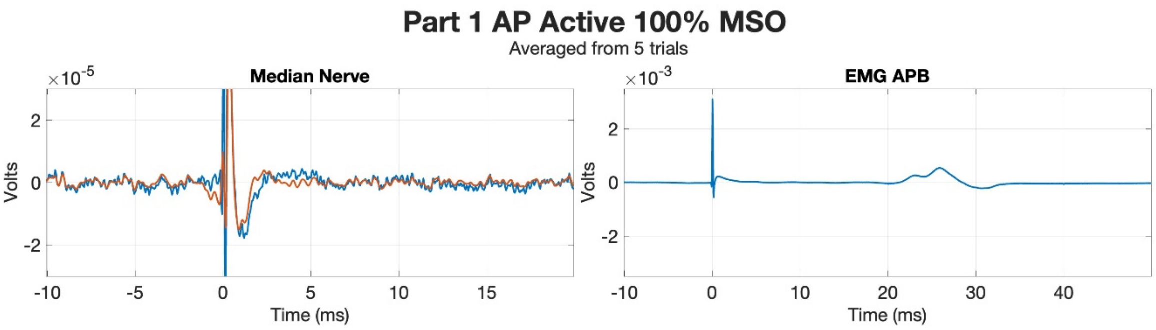
Part 1 AP Active 100% MSO
Averaged from 5 trials
Median Nerve
EMG APB
Volts
Time (ms)

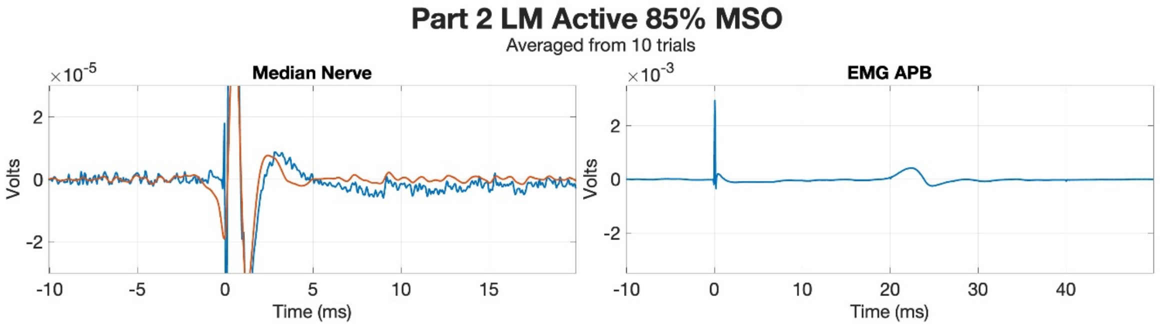
Part 2 LM Active 85% MSO
Averaged from 10 trials
Median Nerve
EMG APB
×10^-5
×10^-3
Volts
Time (ms)

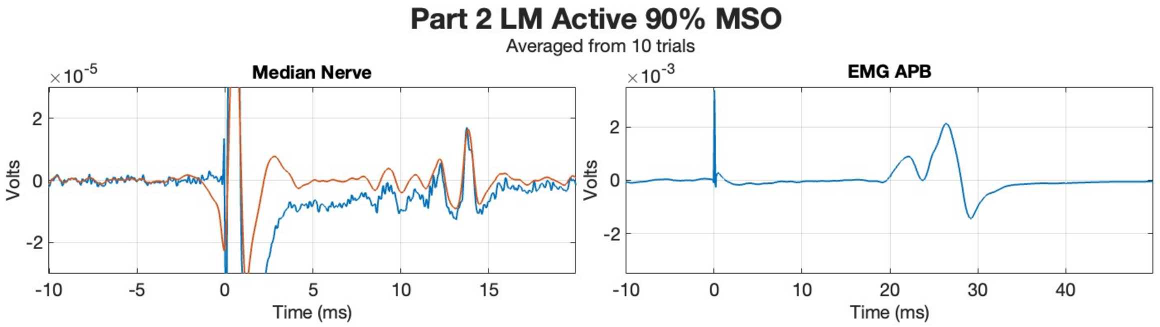
Part 2 LM Active 90% MSO
Averaged from 10 trials
Median Nerve
EMG APB
×10^-5
×10^-3
Volts
Time (ms)

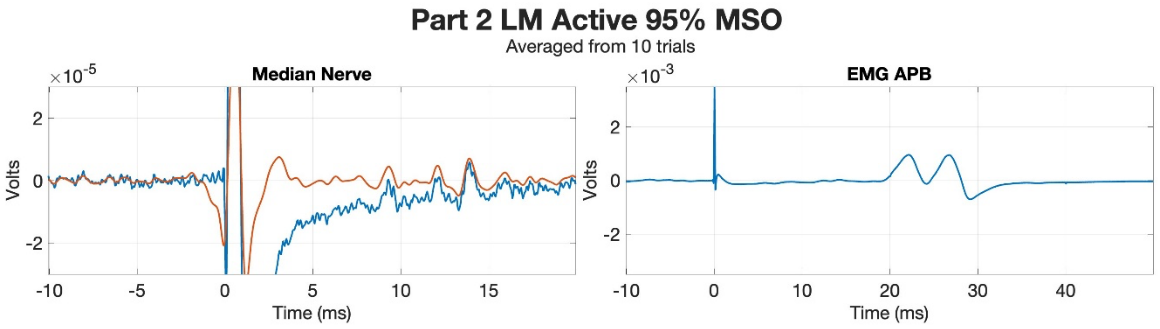
Part 2 LM Active 95% MSO
Averaged from 10 trials
Median Nerve
EMG APB
×10^-5
×10^-3
Volts
Time (ms)

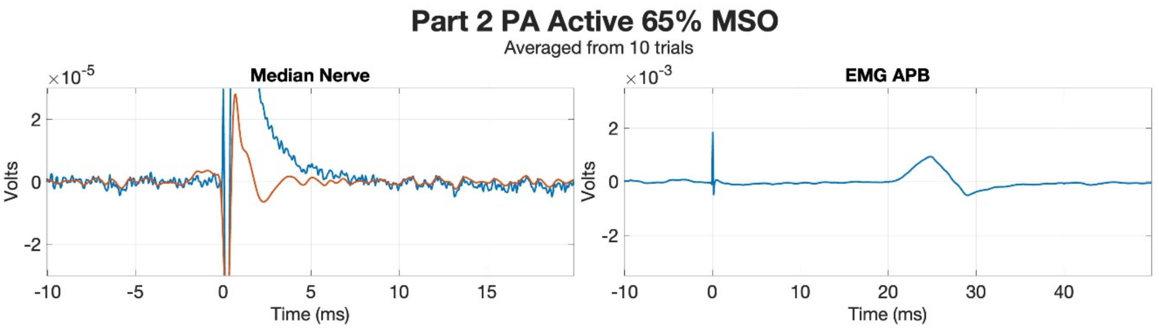
Part 2 PA Active 65% MSO
Averaged from 10 trials
Median Nerve
EMG APB
×10^-5
×10^-3
Volts
Time (ms)

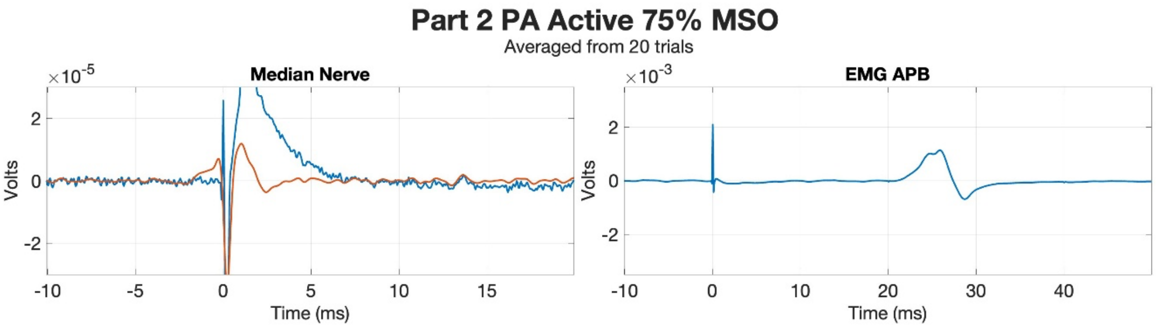
Part 2 PA Active 75% MSO
Averaged from 20 trials
Median Nerve
EMG APB
×10^-5
×10^-3
Volts
Time (ms)

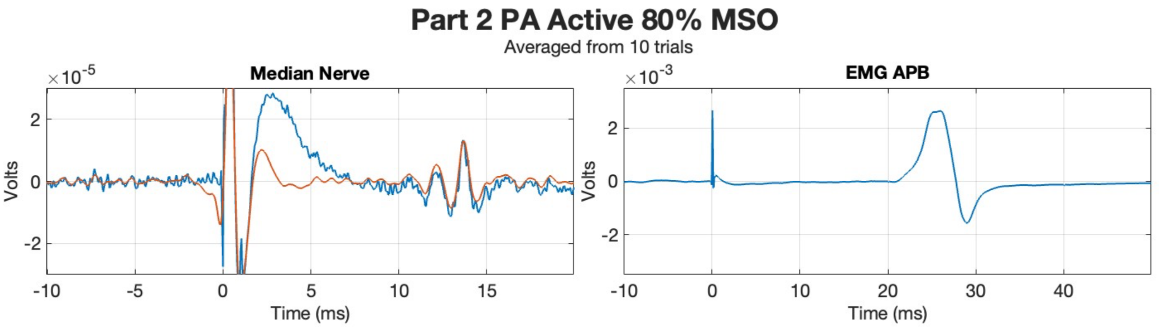
Part 2 PA Active 80% MSO
Averaged from 10 trials
Median Nerve
EMG APB
×10^-5
×10^-3
Volts
Time (ms)

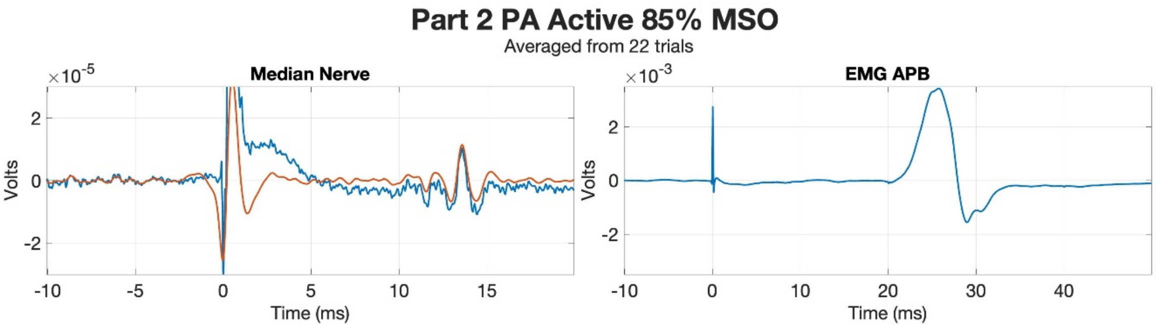
Part 2 PA Active 85% MSO
Averaged from 22 trials
Median Nerve
EMG APB
×10^-5
×10^-3
Volts
Time (ms)

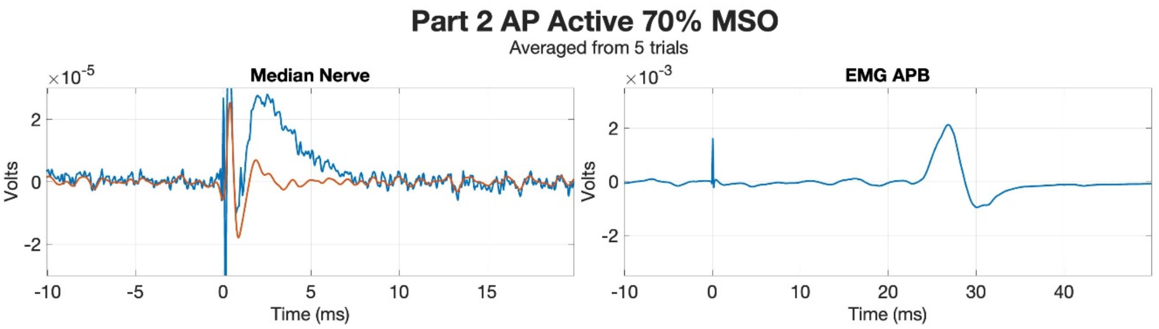
Part 2 AP Active 70% MSO
Averaged from 5 trials
Median Nerve
EMG APB
×10^-5
×10^-3
Volts
Time (ms)

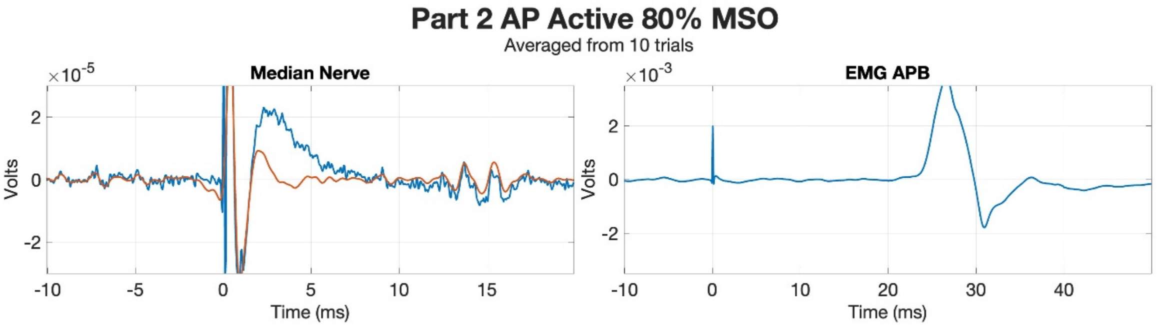
Part 2 AP Active 80% MSO
Averaged from 10 trials
Median Nerve
EMG APB
×10^-5
×10^-3
Volts
Time (ms)

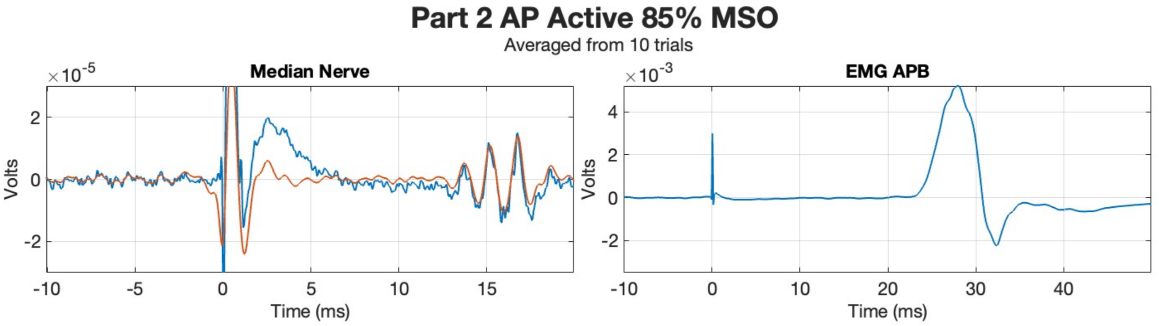
Part 2 AP Active 85% MSO
Averaged from 10 trials
Median Nerve
EMG APB
×10^-5
×10^-3
Volts
Time (ms)

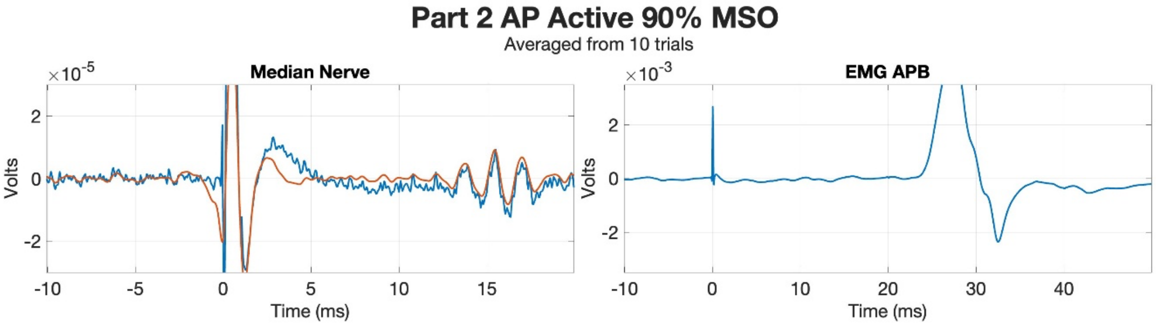
Part 2 AP Active 90% MSO
Averaged from 10 trials
Median Nerve
EMG APB
×10^-5
×10^-3
Volts
Time (ms)

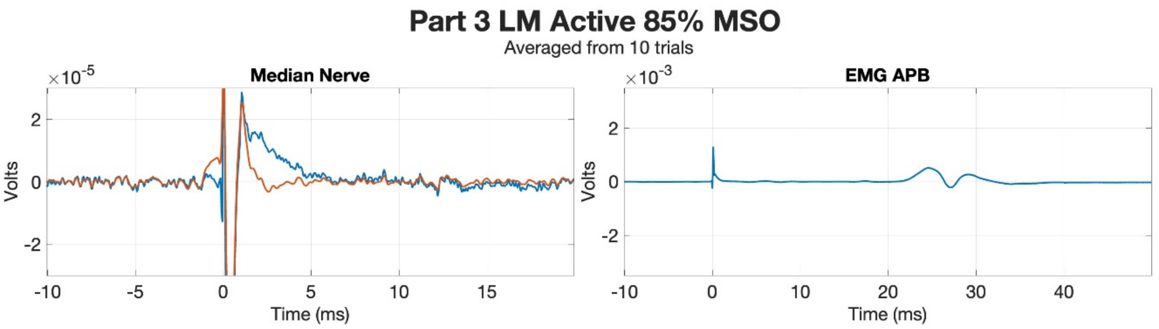
Part 3 LM Active 85% MSO
Averaged from 10 trials
Median Nerve
EMG APB
Volts
Time (ms)

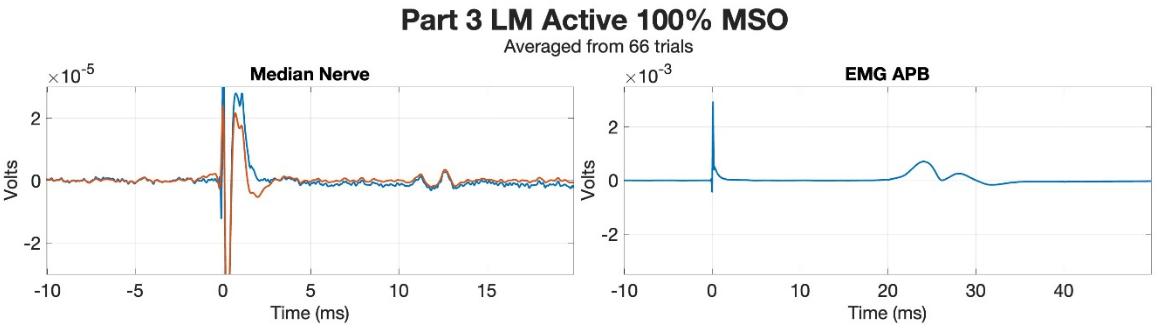
Part 3 LM Active 100% MSO
Averaged from 66 trials
Median Nerve
EMG APB
Volts
Time (ms)

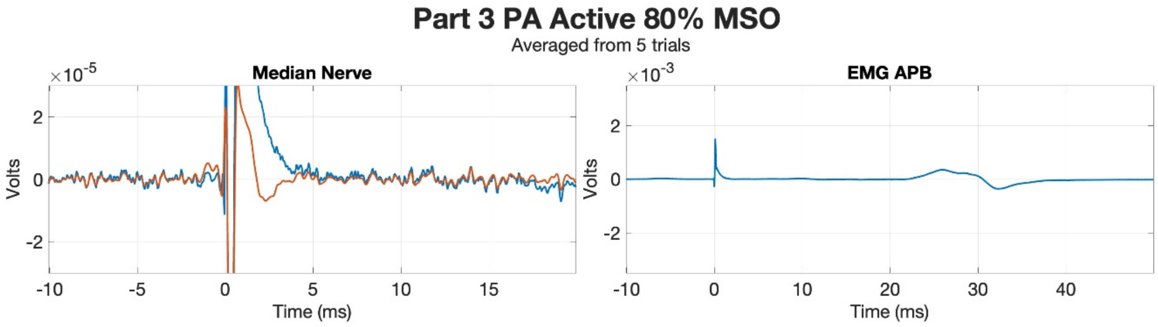
Part 3 PA Active 80% MSO
Averaged from 5 trials
Median Nerve
EMG APB
×10^-5
×10^-3
Volts
Time (ms)

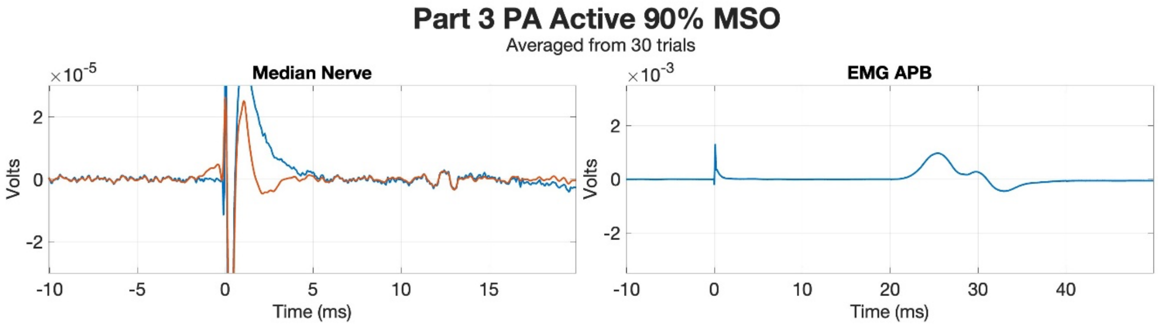
Part 3 PA Active 90% MSO
Averaged from 30 trials
Median Nerve
EMG APB
×10^-5
×10^-3
Volts
Time (ms)

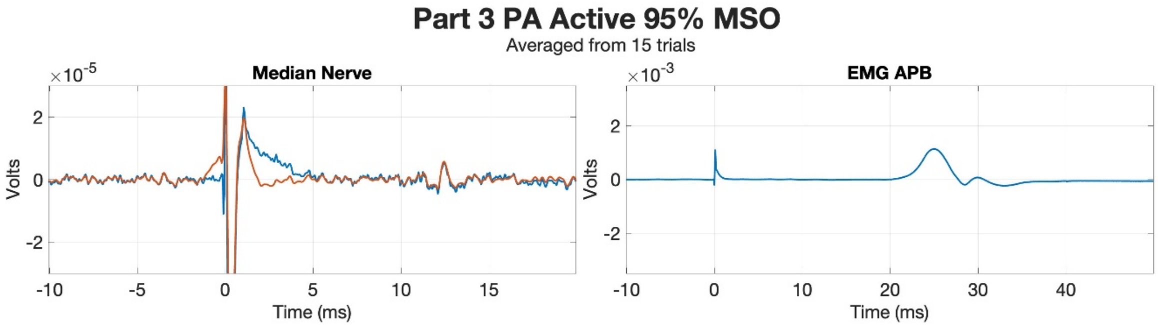
Part 3 PA Active 95% MSO
Averaged from 15 trials
Median Nerve
EMG APB
×10^-5
×10^-3
Volts
Time (ms)

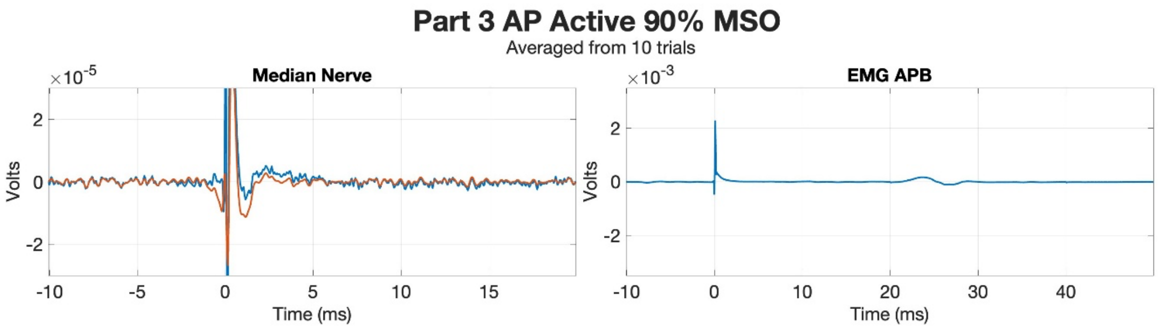
Part 3 AP Active 90% MSO
Averaged from 10 trials
Median Nerve
EMG APB
Volts
Time (ms)

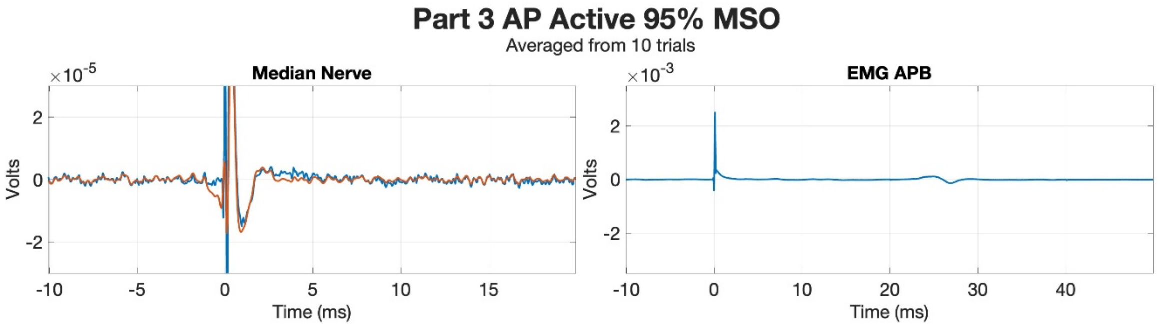
Part 3 AP Active 95% MSO
Averaged from 10 trials
Median Nerve
EMG APB
Volts
Time (ms)

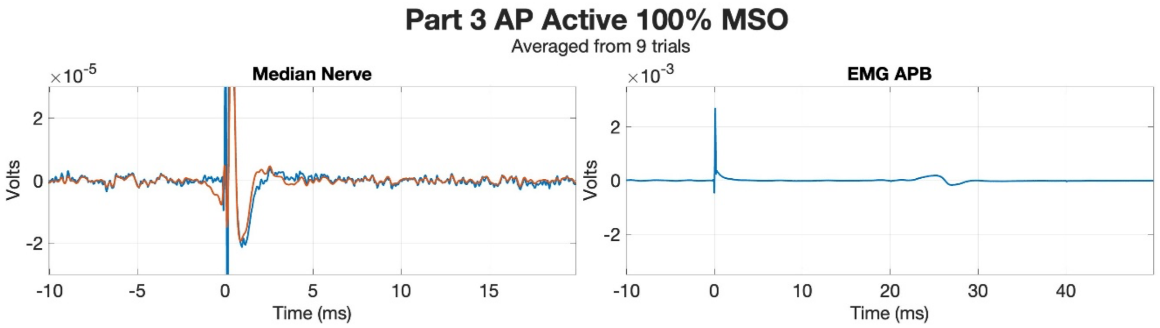
Part 3 AP Active 100% MSO
Averaged from 9 trials
Median Nerve
EMG APB
Volts
Time (ms)

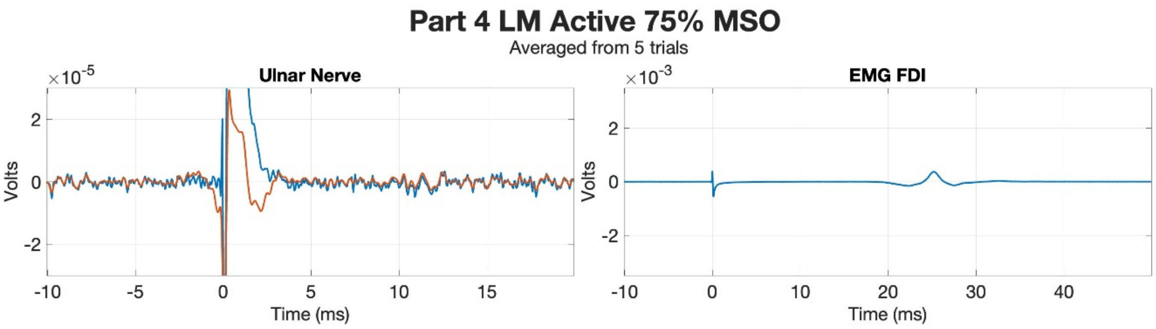
Part 4 LM Active 75% MSO
Averaged from 5 trials
Ulnar Nerve
×10-5
Volts
Time (ms)
EMG FDI
×10-3
Volts
Time (ms)

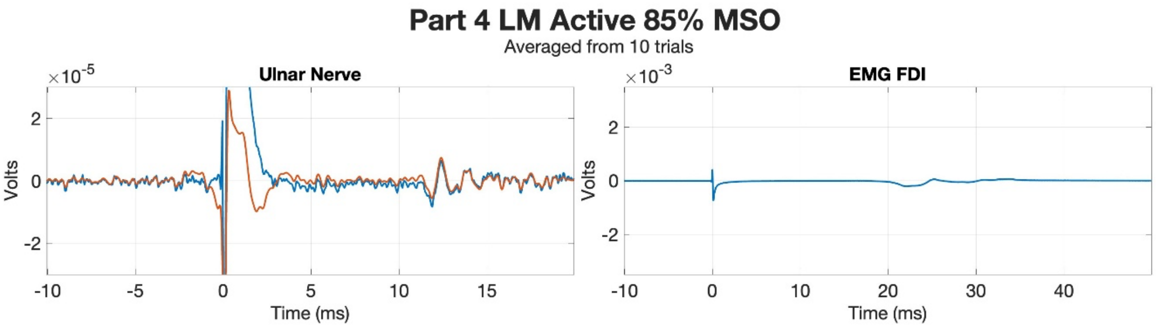
Part 4 LM Active 85% MSO
Averaged from 10 trials
Ulnar Nerve
×10-5
Volts
Time (ms)
EMG FDI
×10-3
Volts
Time (ms)

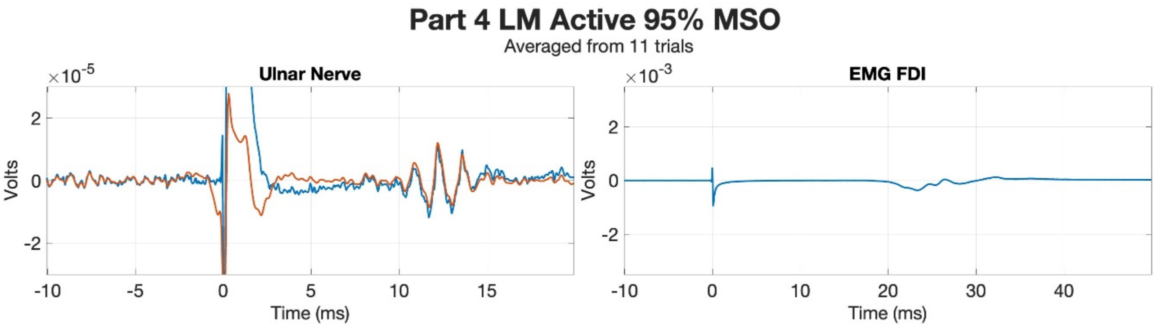
Part 4 LM Active 95% MSO
Averaged from 11 trials
Ulnar Nerve
×10-5
Volts
Time (ms)
EMG FDI
×10-3
Volts
Time (ms)

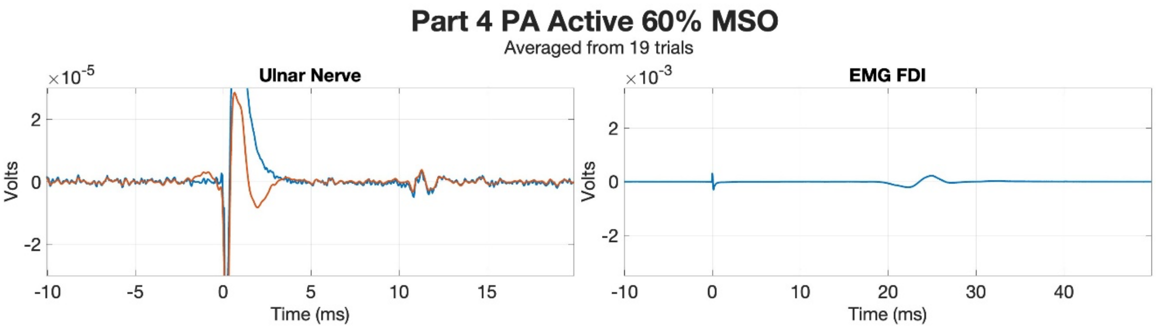
Part 4 PA Active 60% MSO
Averaged from 19 trials
Ulnar Nerve
EMG FDI
×10^-5
×10^-3
Volts
Time (ms)

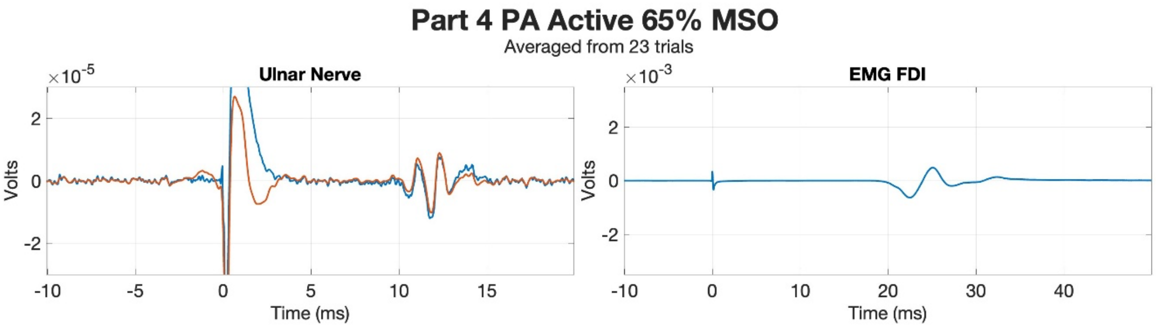
Part 4 PA Active 65% MSO
Averaged from 23 trials
Ulnar Nerve
EMG FDI
×10^-5
×10^-3
Volts
Time (ms)

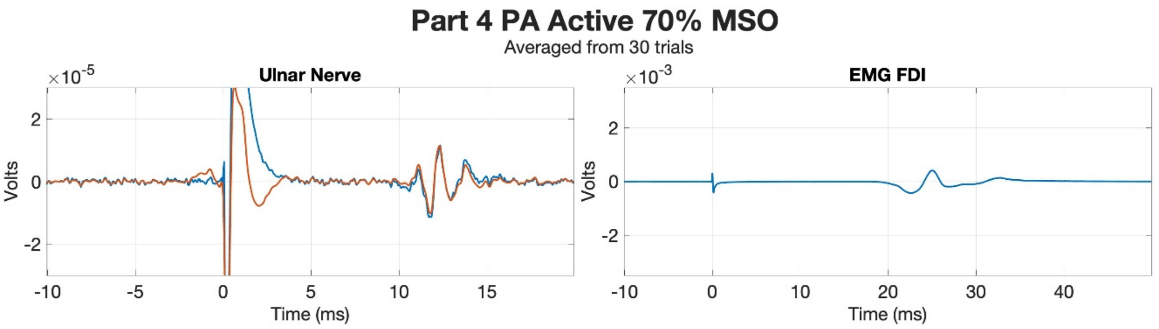
Part 4 PA Active 70% MSO
Averaged from 30 trials
Ulnar Nerve
EMG FDI
×10^-5
×10^-3
Volts
Time (ms)

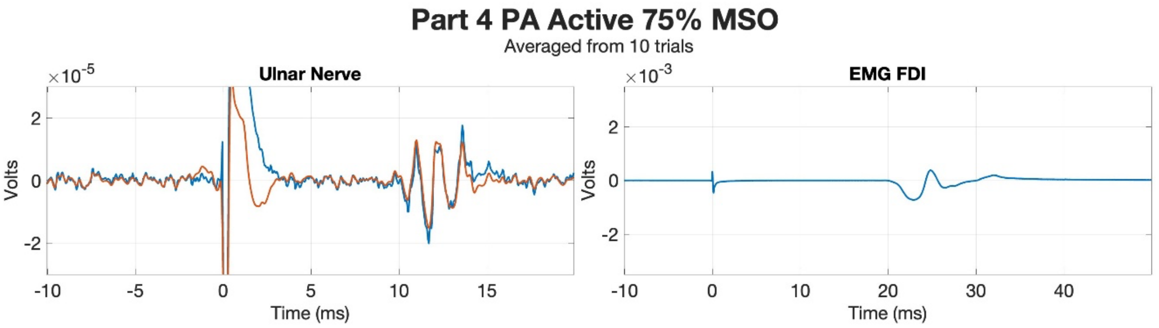
Part 4 PA Active 75% MSO
Averaged from 10 trials
Ulnar Nerve
EMG FDI
×10^-5
×10^-3
Volts
Time (ms)

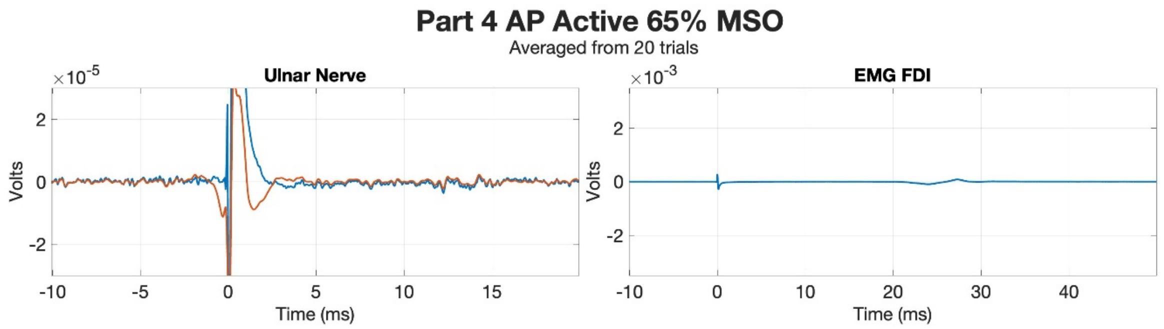
Part 4 AP Active 65% MSO
Averaged from 20 trials
Ulnar Nerve
EMG FDI
×10^-5
×10^-3
Volts
Time (ms)

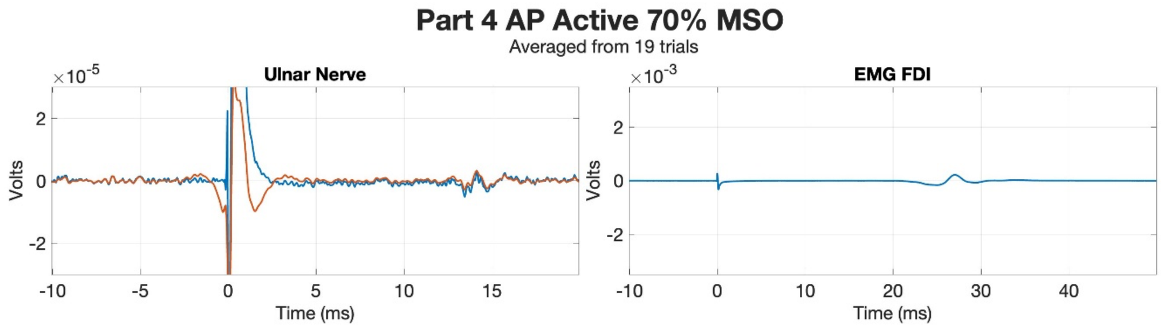
Part 4 AP Active 70% MSO
Averaged from 19 trials
Ulnar Nerve
EMG FDI
×10^-5
×10^-3
Volts
Time (ms)

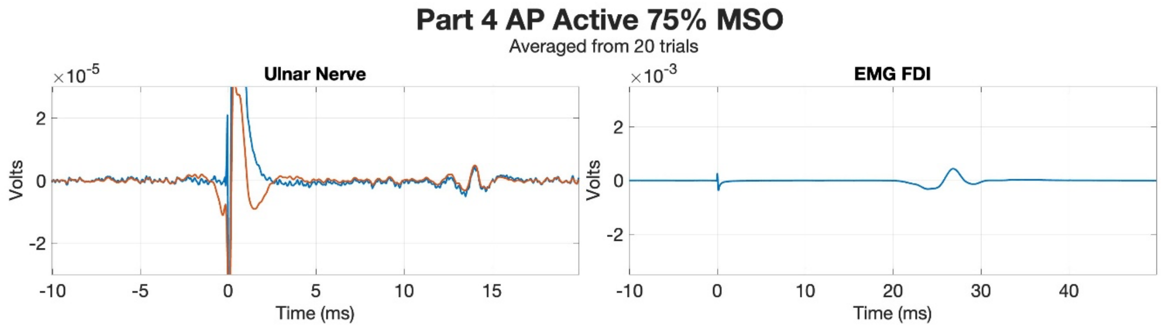
Part 4 AP Active 75% MSO
Averaged from 20 trials
Ulnar Nerve
EMG FDI
×10^-5
×10^-3
Volts
Time (ms)

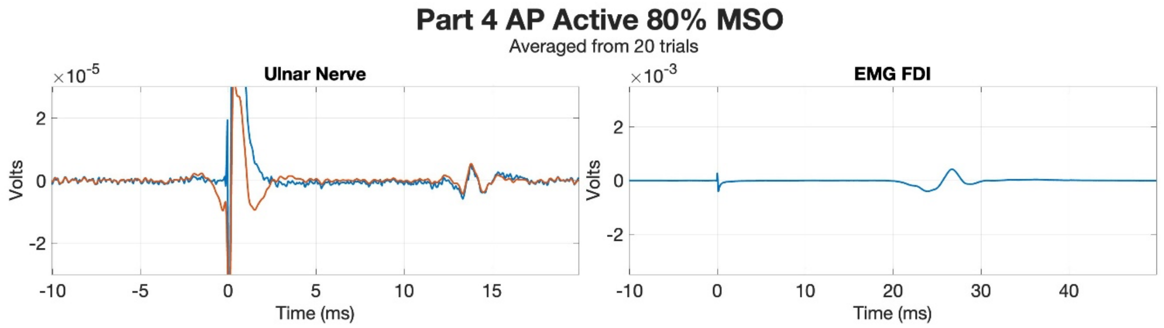
Part 4 AP Active 80% MSO
Averaged from 20 trials
Ulnar Nerve
EMG FDI
×10^-5
×10^-3
Volts
Time (ms)

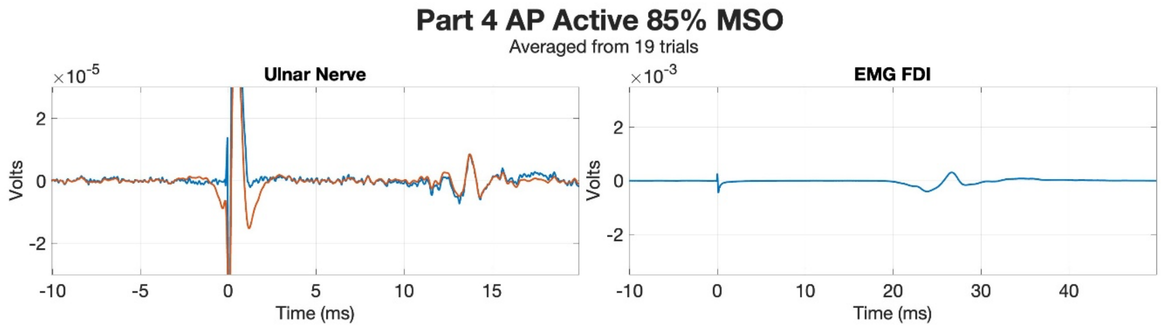
Part 4 AP Active 85% MSO
Averaged from 19 trials
Ulnar Nerve
EMG FDI
×10^-5
×10^-3
Volts
Time (ms)

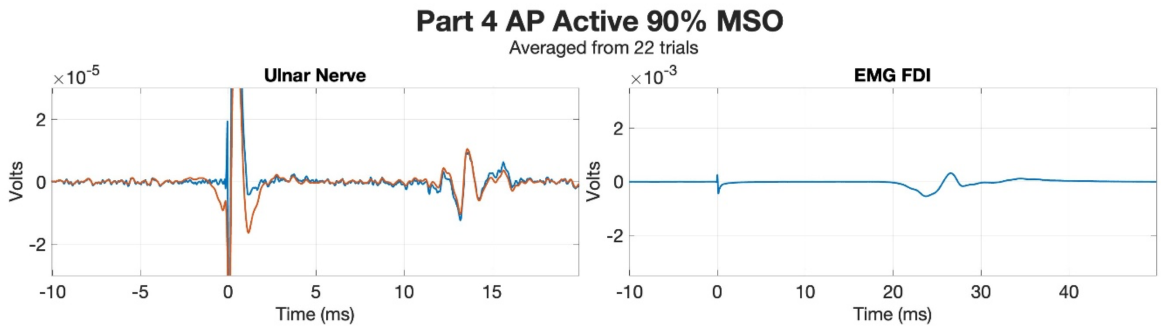
Part 4 AP Active 90% MSO
Averaged from 22 trials
Ulnar Nerve
×10^-5
Volts
Time (ms)
EMG FDI
×10^-3
Volts
Time (ms)

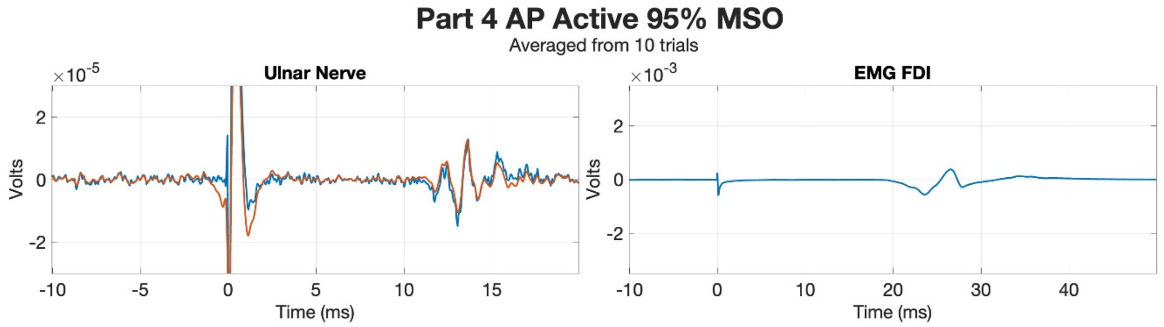
Part 4 AP Active 95% MSO
Averaged from 10 trials
Ulnar Nerve
×10^-5
Volts
Time (ms)
EMG FDI
×10^-3
Volts
Time (ms)

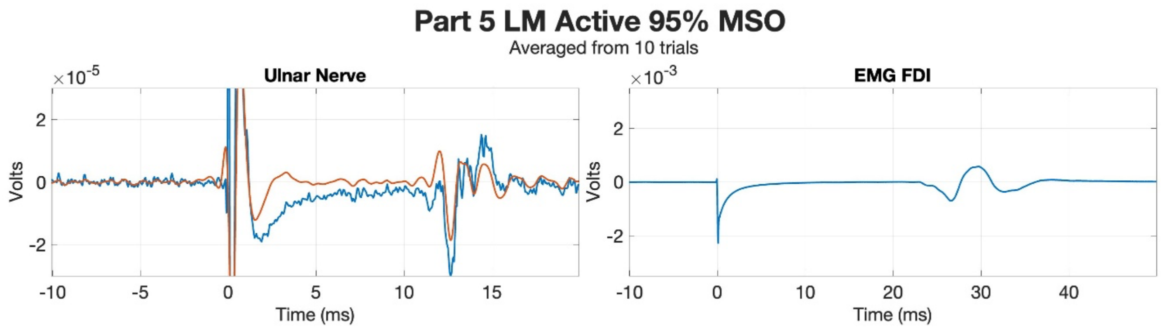
Part 5 LM Active 95% MSO
Averaged from 10 trials
Ulnar Nerve
EMG FDI
Volts
Time (ms)

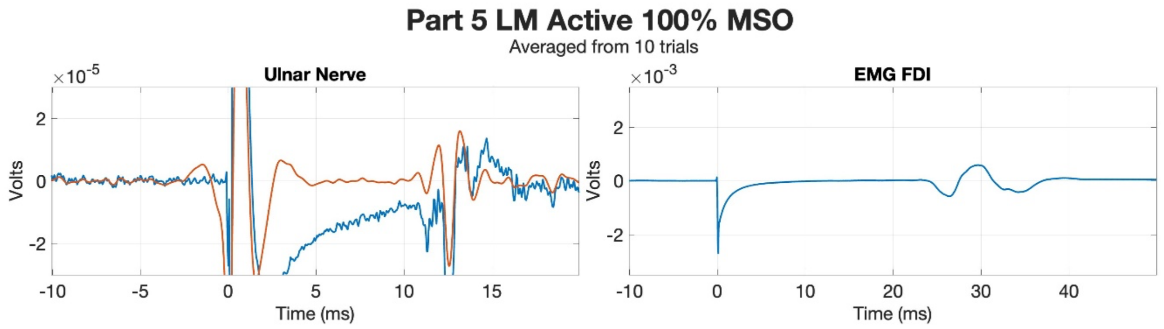
Part 5 LM Active 100% MSO
Averaged from 10 trials
Ulnar Nerve
EMG FDI
Volts
Time (ms)

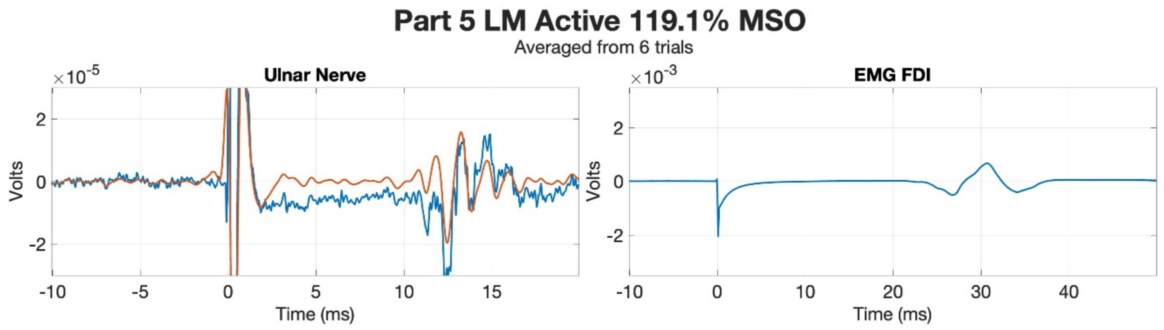
Part 5 LM Active 119.1% MSO
Averaged from 6 trials
Ulnar Nerve
EMG FDI
Volts
Time (ms)

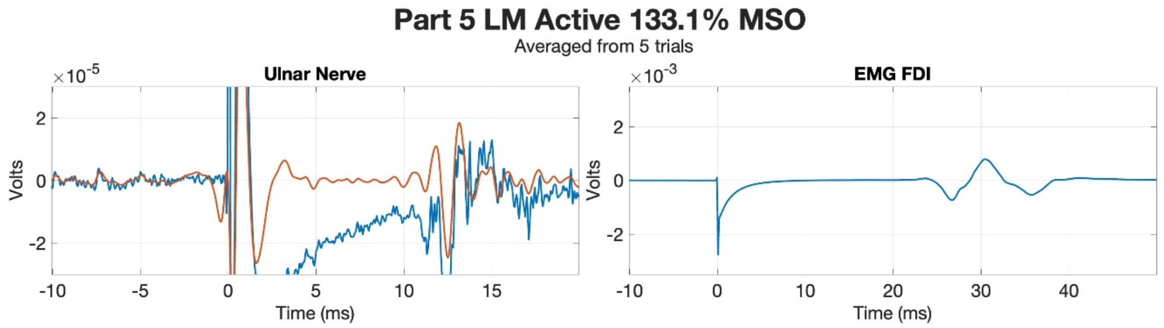
Part 5 LM Active 133.1% MSO
Averaged from 5 trials
Ulnar Nerve
EMG FDI
Volts
Time (ms)

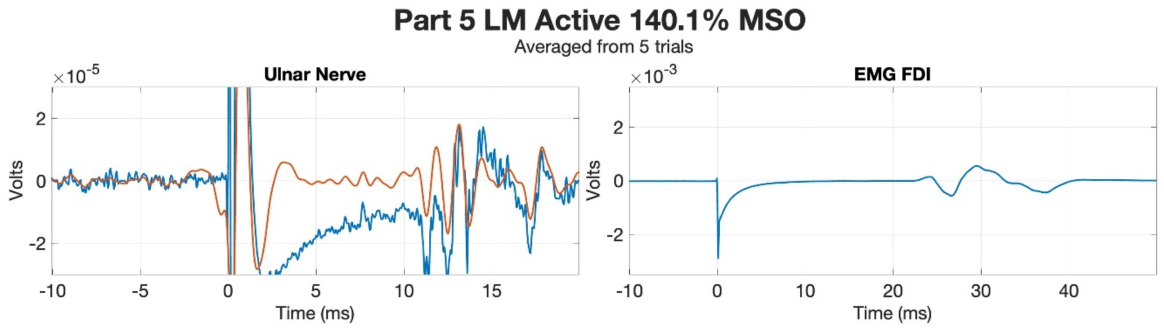
Part 5 LM Active 140.1% MSO
Averaged from 5 trials
Ulnar Nerve
EMG FDI
Volts
Time (ms)

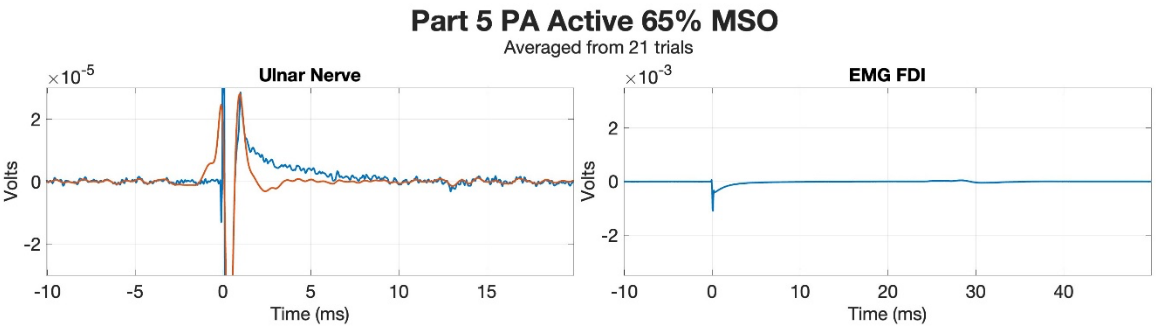
Part 5 PA Active 65% MSO
Averaged from 21 trials
Ulnar Nerve
EMG FDI
×10-5
×10-3
Volts
Time (ms)

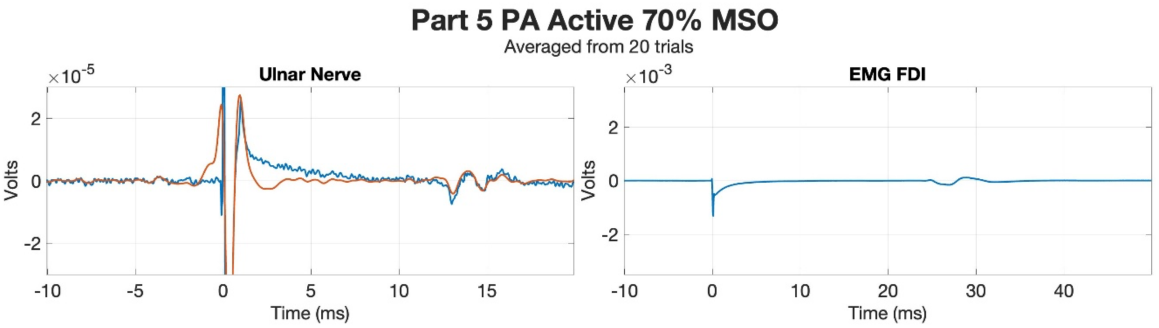
Part 5 PA Active 70% MSO
Averaged from 20 trials
Ulnar Nerve
EMG FDI
×10-5
×10-3
Volts
Time (ms)

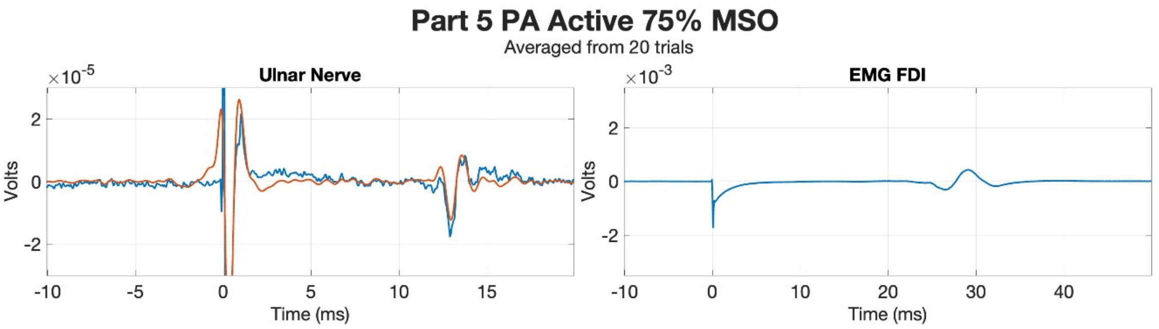
Part 5 PA Active 75% MSO
Averaged from 20 trials
Ulnar Nerve
EMG FDI
×10-5
×10-3
Volts
Time (ms)

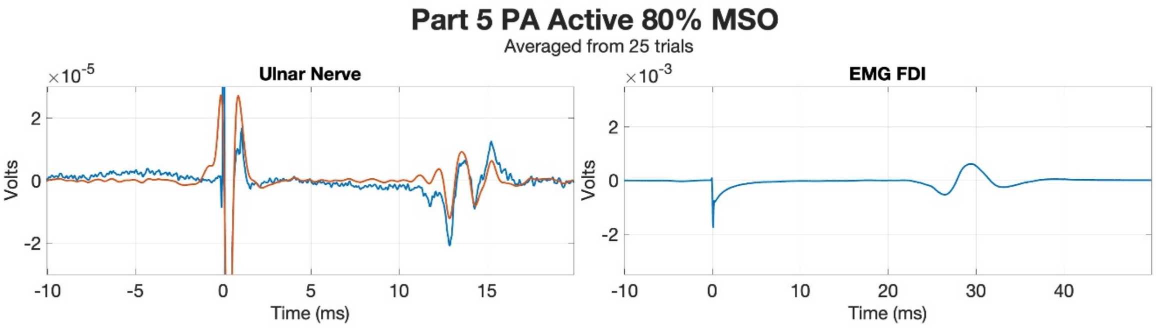
Part 5 PA Active 80% MSO
Averaged from 25 trials
Ulnar Nerve
EMG FDI
×10-5
×10-3
Volts
Time (ms)

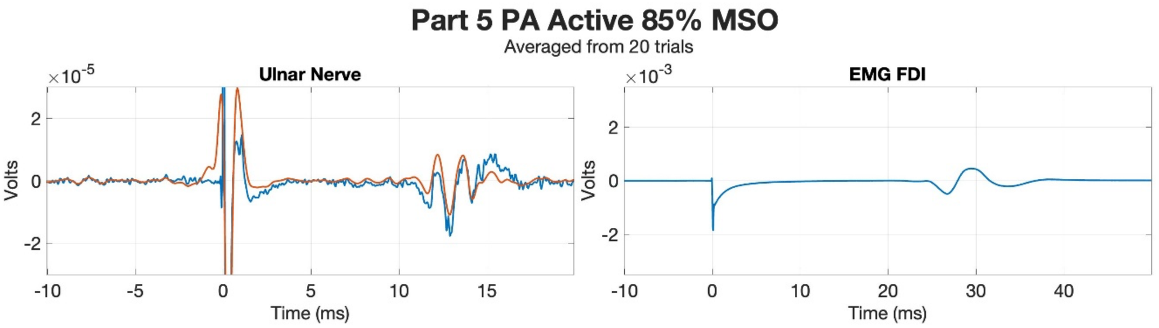
Part 5 PA Active 85% MSO
Averaged from 20 trials
Ulnar Nerve
EMG FDI
×10-5
×10-3
Volts
Time (ms)

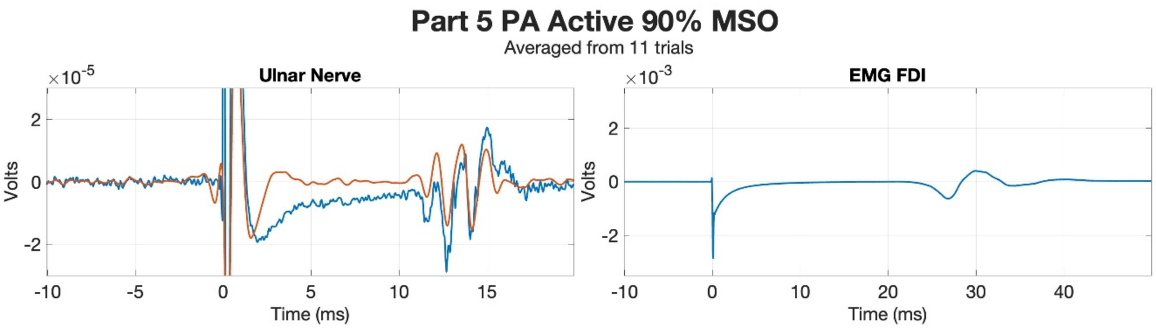
Part 5 PA Active 90% MSO
Averaged from 11 trials
Ulnar Nerve
×10^-5
Volts
Time (ms)
EMG FDI
×10^-3
Volts
Time (ms)

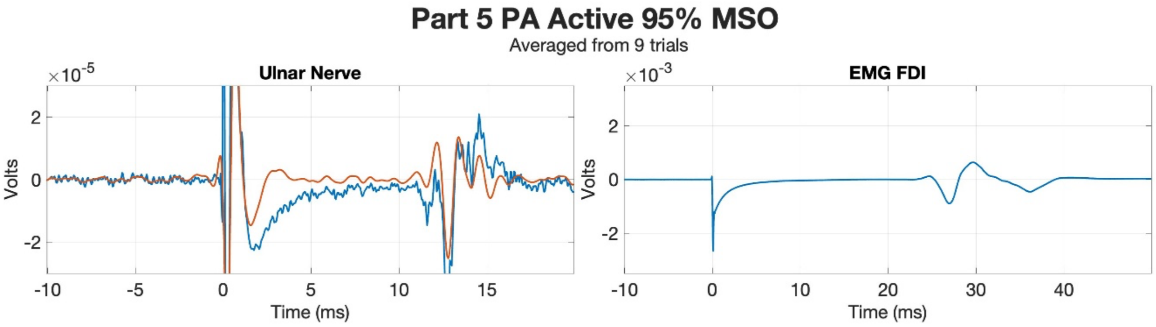
Part 5 PA Active 95% MSO
Averaged from 9 trials
Ulnar Nerve
×10^-5
Volts
Time (ms)
EMG FDI
×10^-3
Volts
Time (ms)

**Part 5 AP Active 65% MSO**
Averaged from 10 trials

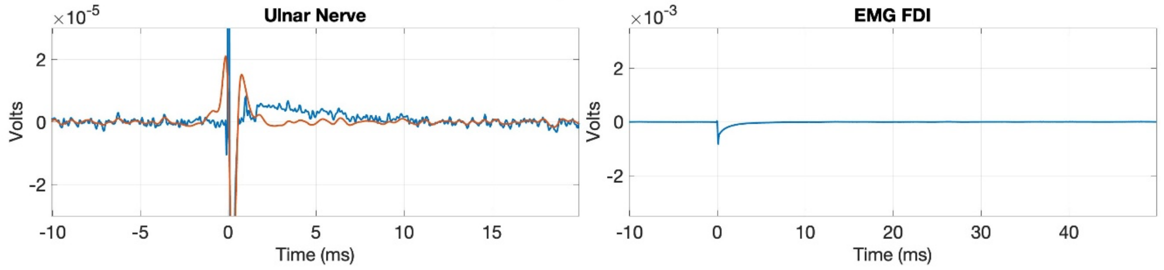


**Part 5 AP Active 70% MSO**
Averaged from 15 trials

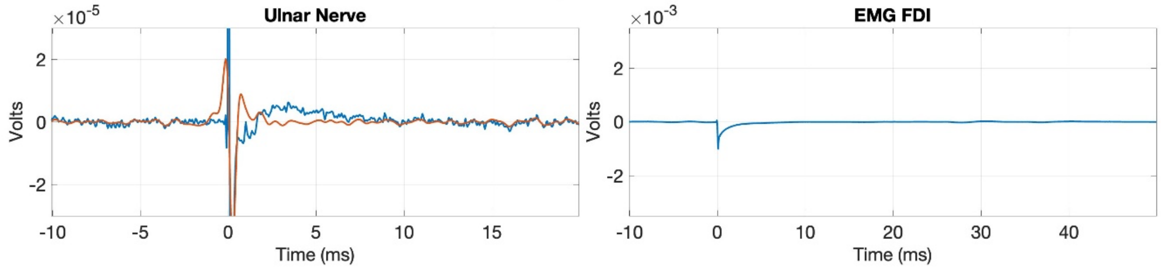


**Part 5 AP Active 75% MSO**
Averaged from 20 trials

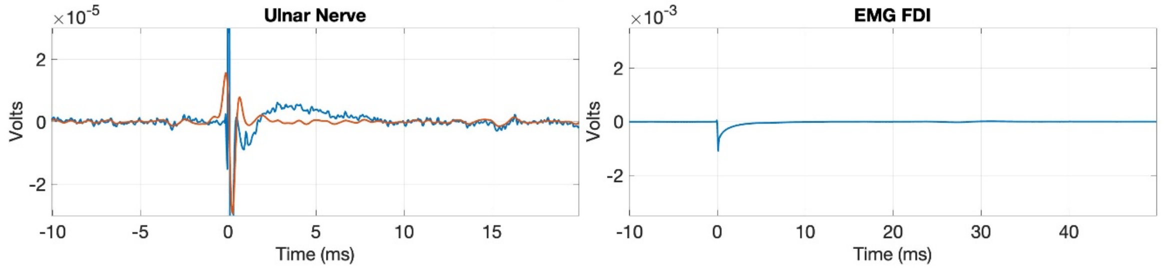


**Part 5 AP Active 80% MSO**
Averaged from 20 trials

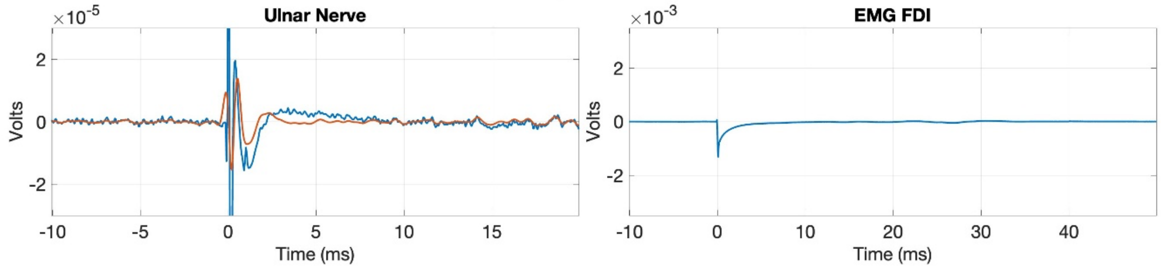


**Part 5 AP Active 85% MSO**
Averaged from 20 trials

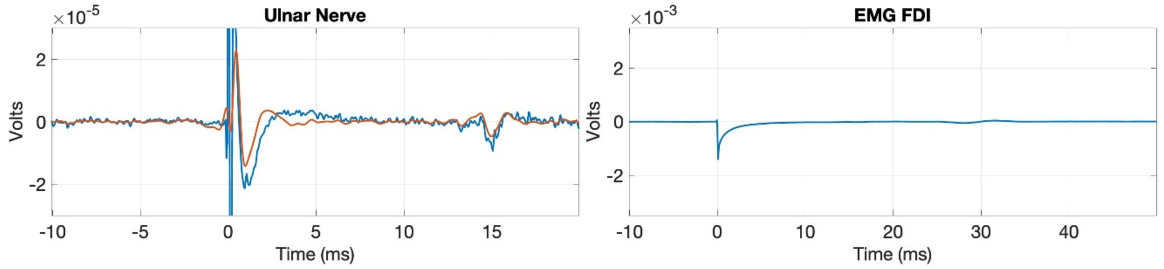

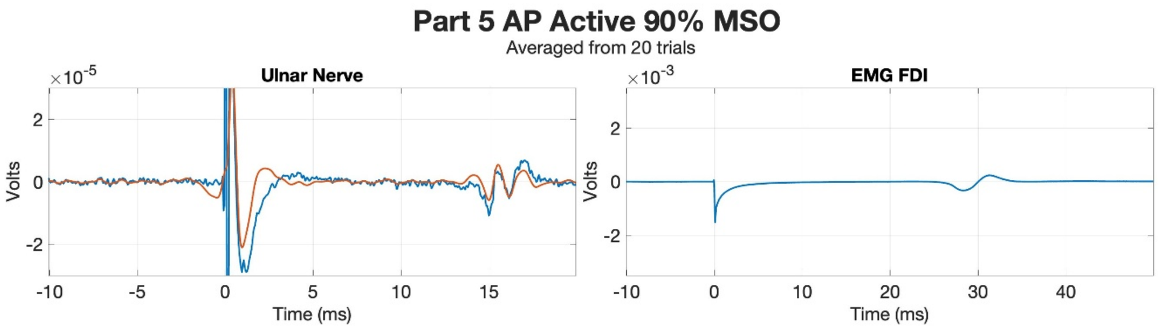
Part 5 AP Active 90% MSO
Averaged from 20 trials
Ulnar Nerve
EMG FDI
×10^-5
×10^-3
Volts
Time (ms)
-10 -5 0 5 10 15
-10 0 10 20 30 40
2 0 -2

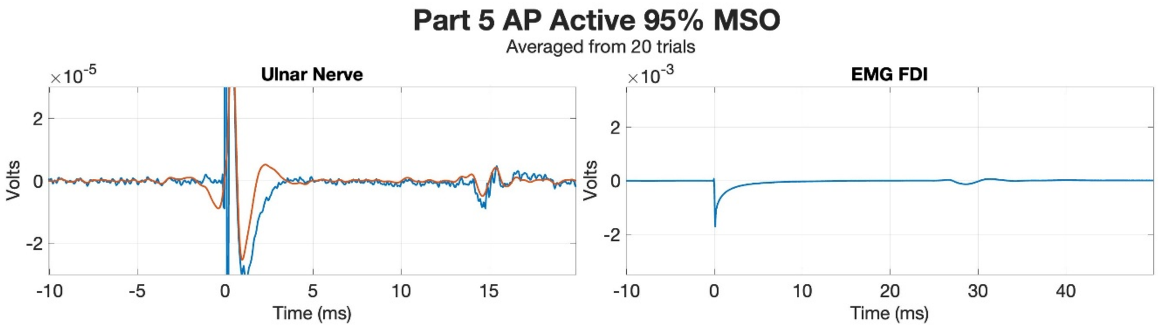
Part 5 AP Active 95% MSO
Averaged from 20 trials
Ulnar Nerve
EMG FDI
×10^-5
×10^-3
Volts
Time (ms)
-10 -5 0 5 10 15
-10 0 10 20 30 40
2 0 -2

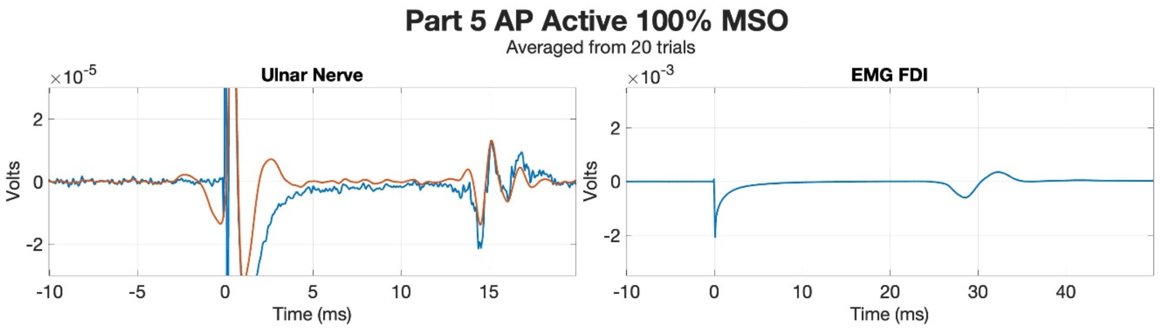
Part 5 AP Active 100% MSO
Averaged from 20 trials
Ulnar Nerve
EMG FDI
×10^-5
×10^-3
Volts
Time (ms)
-10 -5 0 5 10 15
-10 0 10 20 30 40
2 0 -2

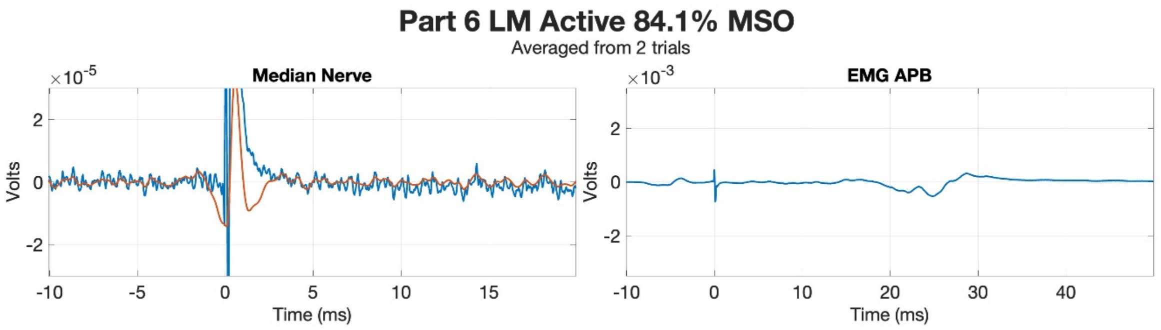
Part 6 LM Active 84.1% MSO
Averaged from 2 trials
Median Nerve
EMG APB
$\times 10^{-5}$
$\times 10^{-3}$
Volts
Time (ms)

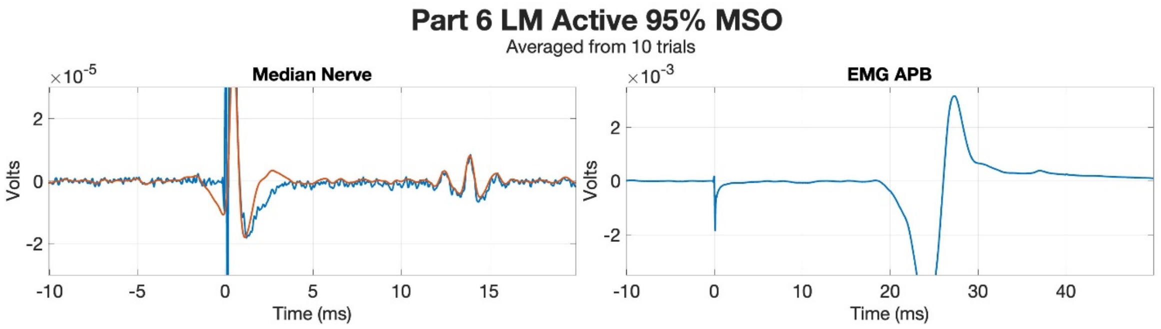
Part 6 LM Active 95% MSO
Averaged from 10 trials
Median Nerve
EMG APB
$\times 10^{-5}$
$\times 10^{-3}$
Volts
Time (ms)

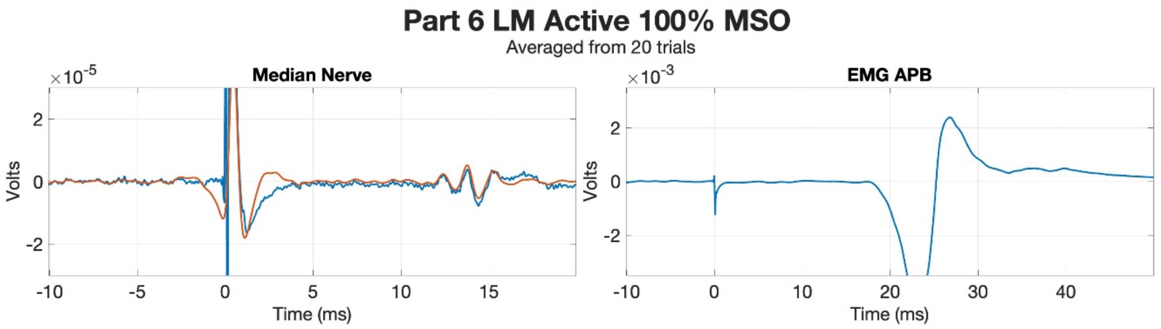
Part 6 LM Active 100% MSO
Averaged from 20 trials
Median Nerve
EMG APB
$\times 10^{-5}$
$\times 10^{-3}$
Volts
Time (ms)

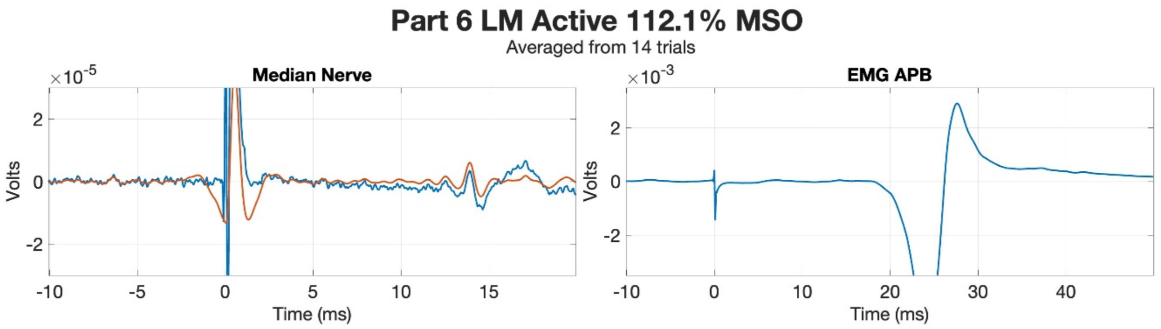
Part 6 LM Active 112.1% MSO
Averaged from 14 trials
Median Nerve
EMG APB
$\times 10^{-5}$
$\times 10^{-3}$
Volts
Time (ms)

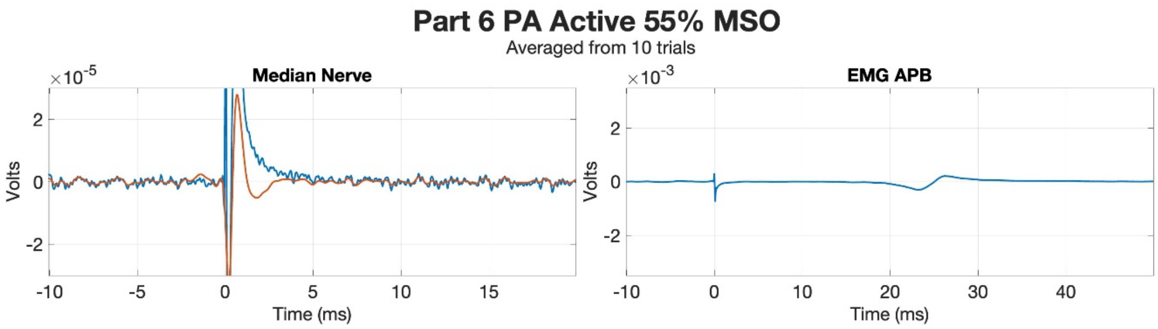
Part 6 PA Active 55% MSO
Averaged from 10 trials
Median Nerve
×10-5
Volts
Time (ms)
EMG APB
×10-3
Volts
Time (ms)

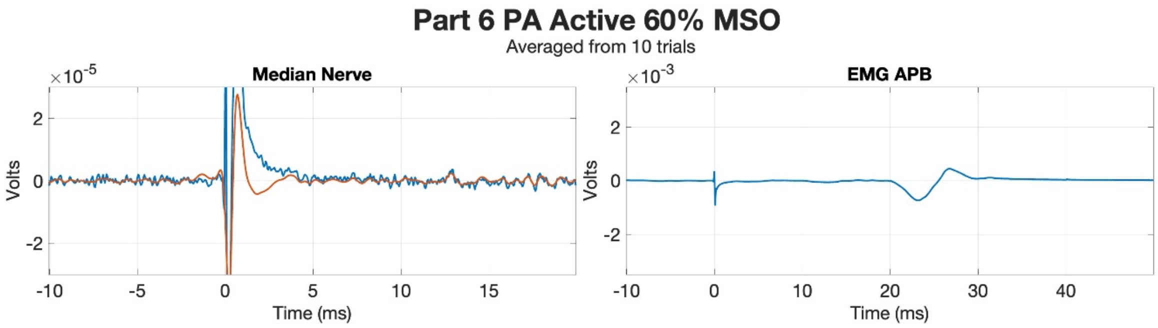
Part 6 PA Active 60% MSO
Averaged from 10 trials
Median Nerve
×10-5
Volts
Time (ms)
EMG APB
×10-3
Volts
Time (ms)

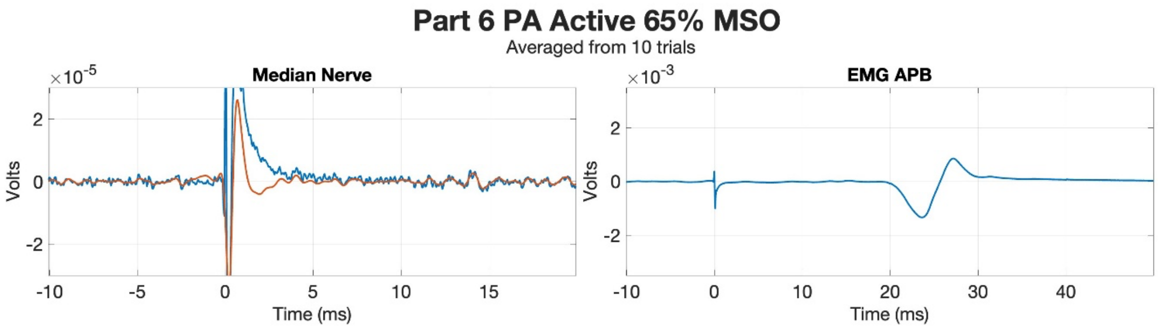
Part 6 PA Active 65% MSO
Averaged from 10 trials
Median Nerve
×10-5
Volts
Time (ms)
EMG APB
×10-3
Volts
Time (ms)

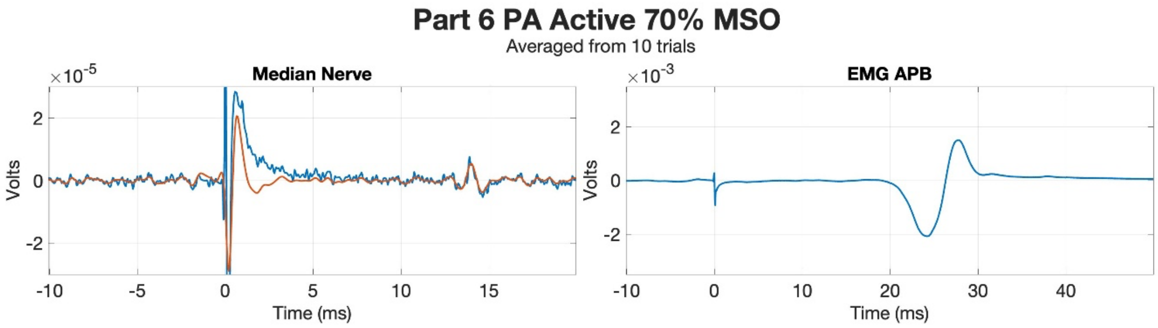
Part 6 PA Active 70% MSO
Averaged from 10 trials
Median Nerve
×10-5
Volts
Time (ms)
EMG APB
×10-3
Volts
Time (ms)

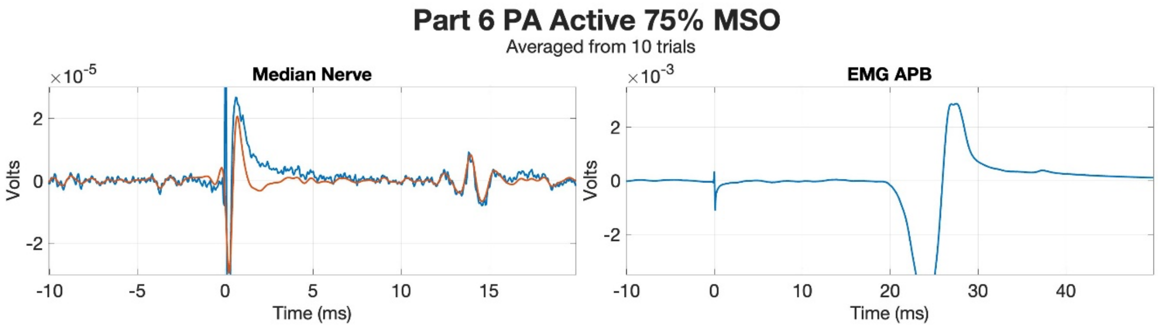
Part 6 PA Active 75% MSO
Averaged from 10 trials
Median Nerve
×10-5
Volts
Time (ms)
EMG APB
×10-3
Volts
Time (ms)

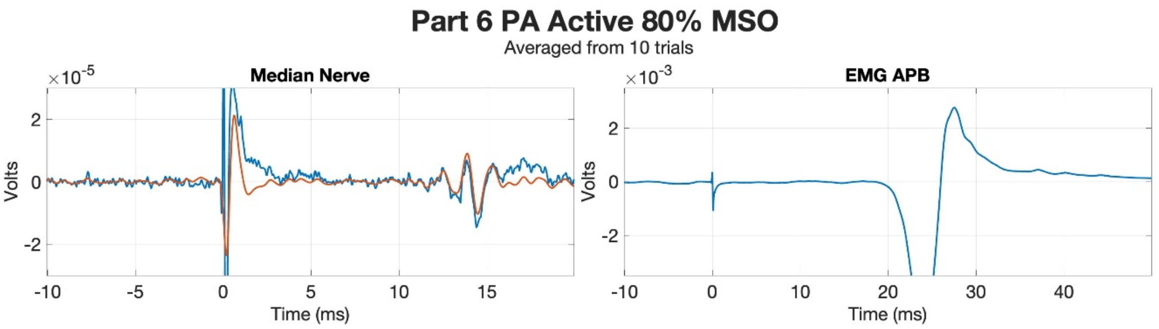
Part 6 PA Active 80% MSO
Averaged from 10 trials
Median Nerve
EMG APB
Volts
Time (ms)

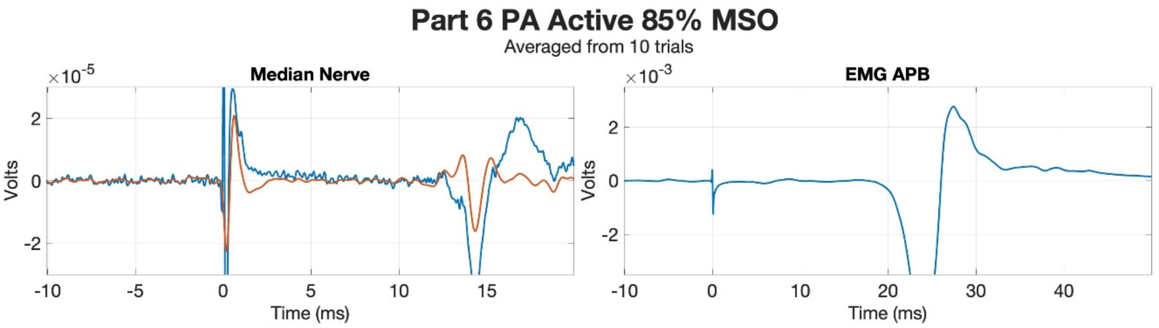
Part 6 PA Active 85% MSO
Averaged from 10 trials
Median Nerve
EMG APB
Volts
Time (ms)

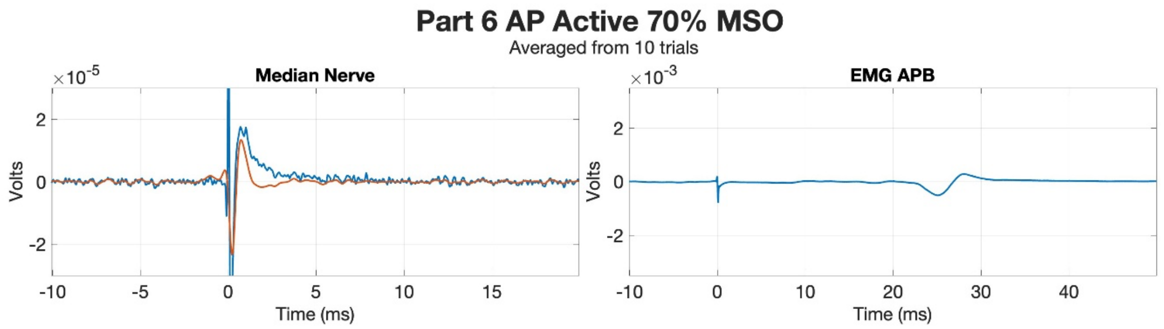
Part 6 AP Active 70% MSO
Averaged from 10 trials
Median Nerve
EMG APB
×10^-5
×10^-3
Volts
Time (ms)

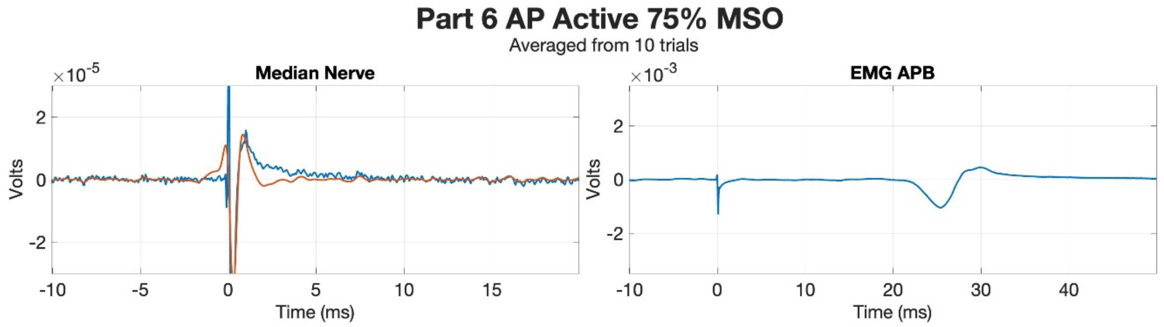
Part 6 AP Active 75% MSO
Averaged from 10 trials
Median Nerve
EMG APB
×10^-5
×10^-3
Volts
Time (ms)

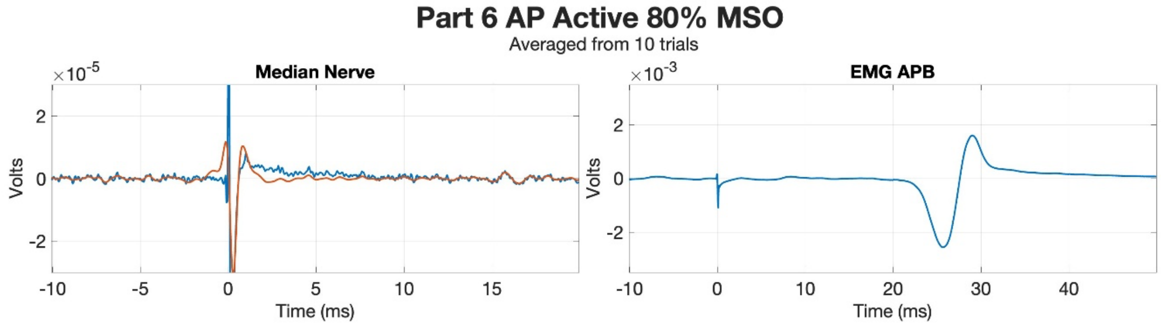
Part 6 AP Active 80% MSO
Averaged from 10 trials
Median Nerve
EMG APB
×10^-5
×10^-3
Volts
Time (ms)

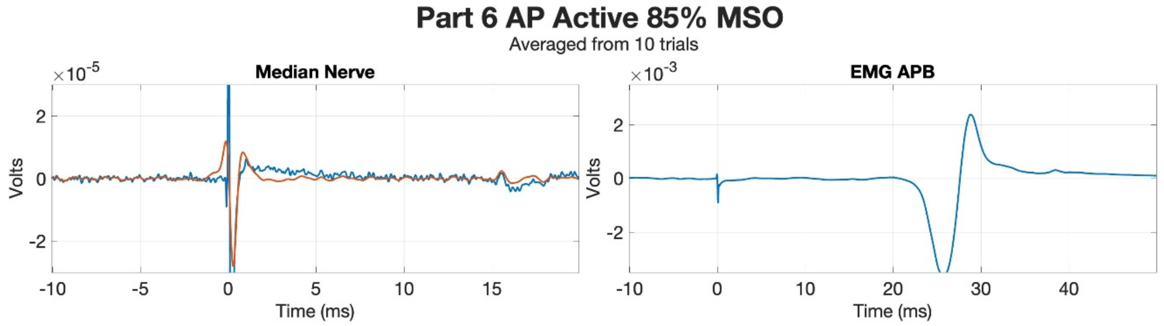
Part 6 AP Active 85% MSO
Averaged from 10 trials
Median Nerve
EMG APB
×10^-5
×10^-3
Volts
Time (ms)

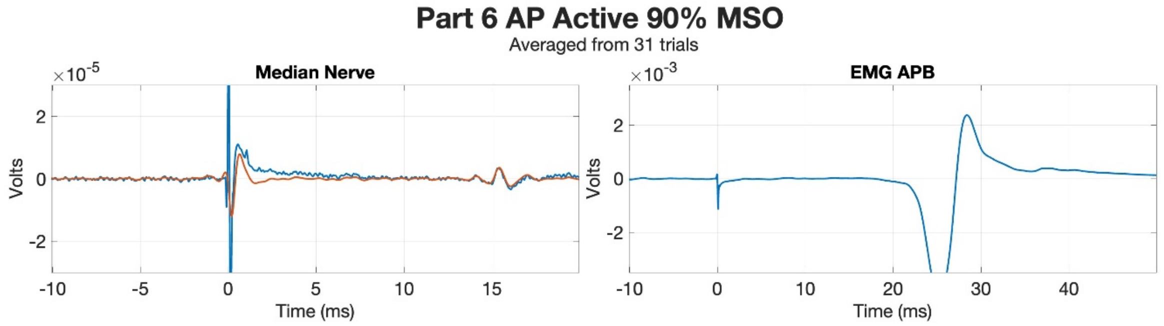
Part 6 AP Active 90% MSO
Averaged from 31 trials
Median Nerve
EMG APB
×10^-5
×10^-3
Volts
Time (ms)

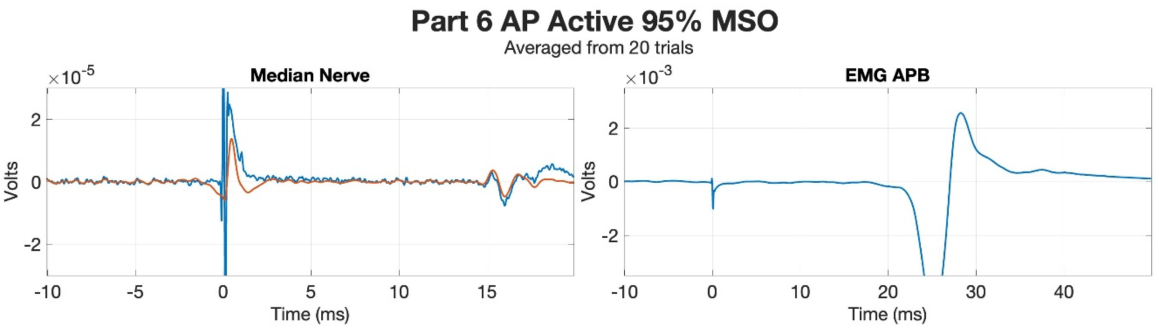
Part 6 AP Active 95% MSO
Averaged from 20 trials
Median Nerve
EMG APB
×10^-5
×10^-3
Volts
Time (ms)

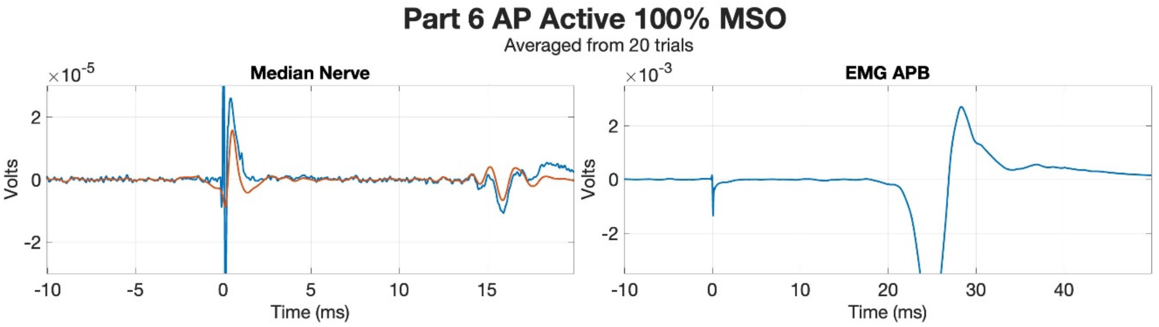
Part 6 AP Active 100% MSO
Averaged from 20 trials
Median Nerve
EMG APB
×10^-5
×10^-3
Volts
Time (ms)

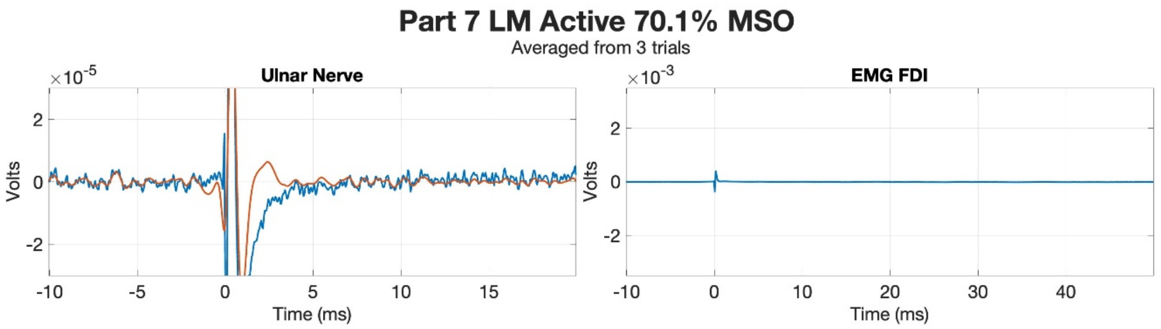
Part 7 LM Active 70.1% MSO
Averaged from 3 trials
Ulnar Nerve
×10^-5
Volts
Time (ms)
EMG FDI
×10^-3
Volts
Time (ms)

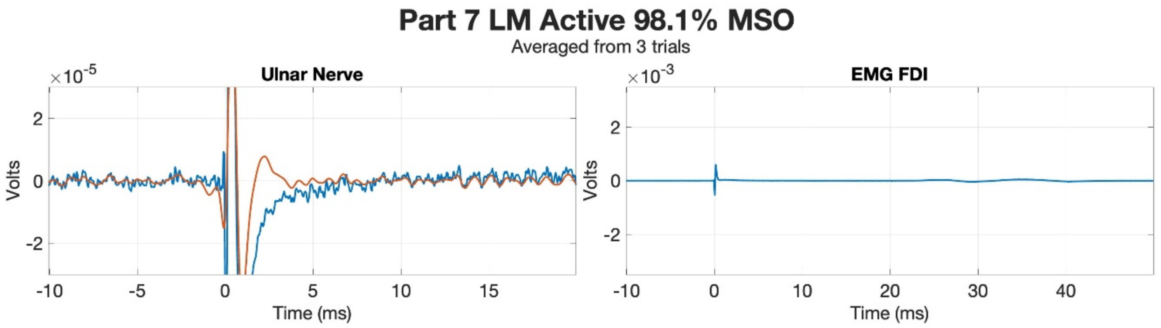
Part 7 LM Active 98.1% MSO
Averaged from 3 trials
Ulnar Nerve
×10^-5
Volts
Time (ms)
EMG FDI
×10^-3
Volts
Time (ms)

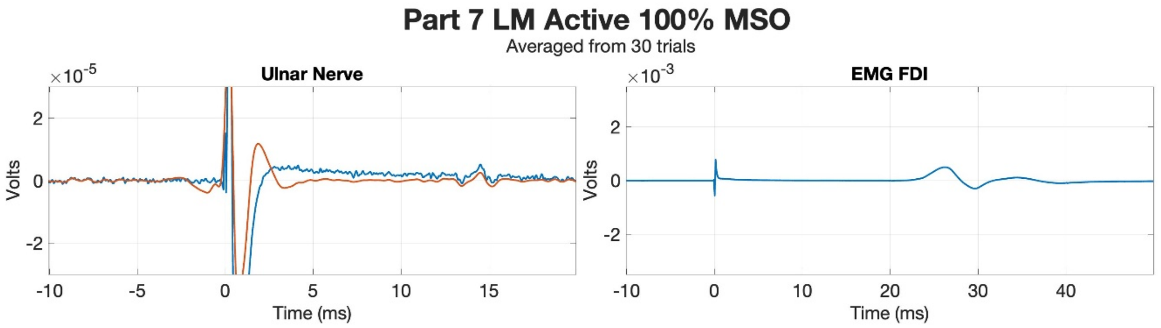
Part 7 LM Active 100% MSO
Averaged from 30 trials
Ulnar Nerve
×10^-5
Volts
Time (ms)
EMG FDI
×10^-3
Volts
Time (ms)

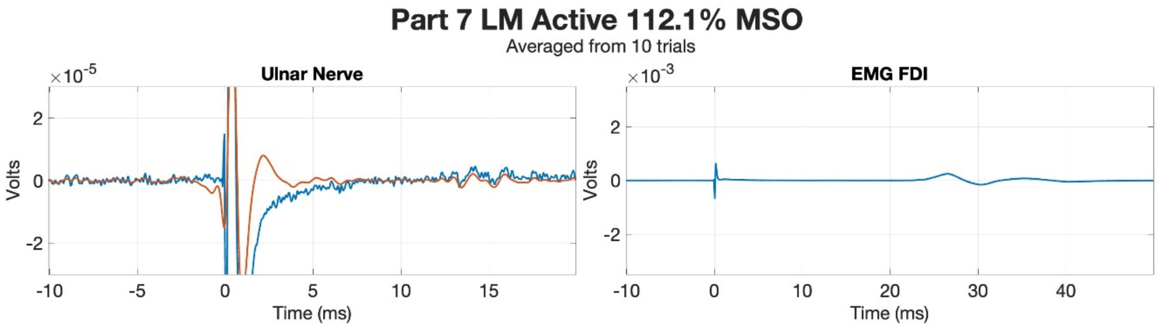
Part 7 LM Active 112.1% MSO
Averaged from 10 trials
Ulnar Nerve
×10^-5
Volts
Time (ms)
EMG FDI
×10^-3
Volts
Time (ms)

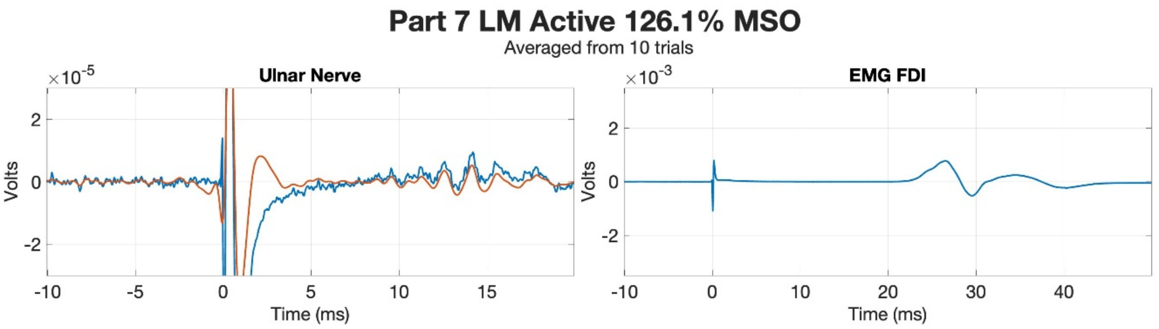
Part 7 LM Active 126.1% MSO
Averaged from 10 trials
Ulnar Nerve
EMG FDI
×10^-5
×10^-3
Volts
Time (ms)

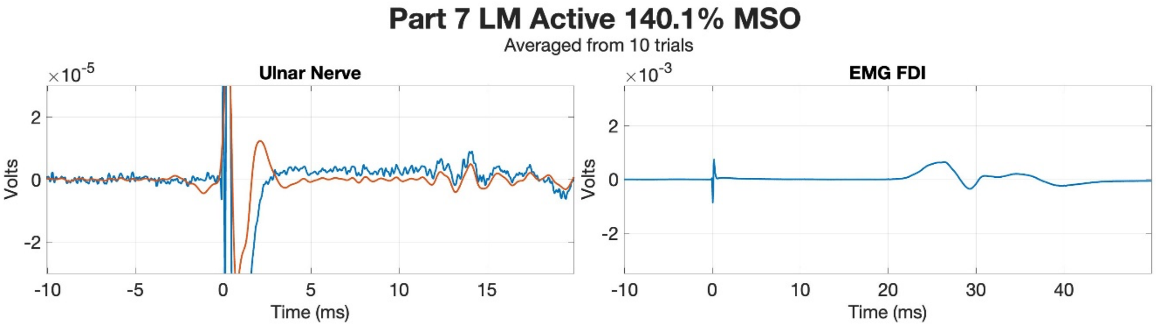
Part 7 LM Active 140.1% MSO
Averaged from 10 trials
Ulnar Nerve
EMG FDI
×10^-5
×10^-3
Volts
Time (ms)

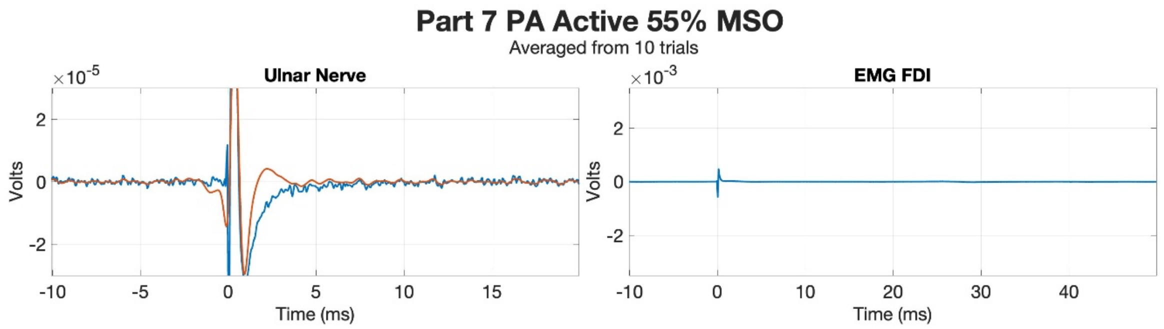
Part 7 PA Active 55% MSO
Averaged from 10 trials
Ulnar Nerve
EMG FDI
Volts
Time (ms)

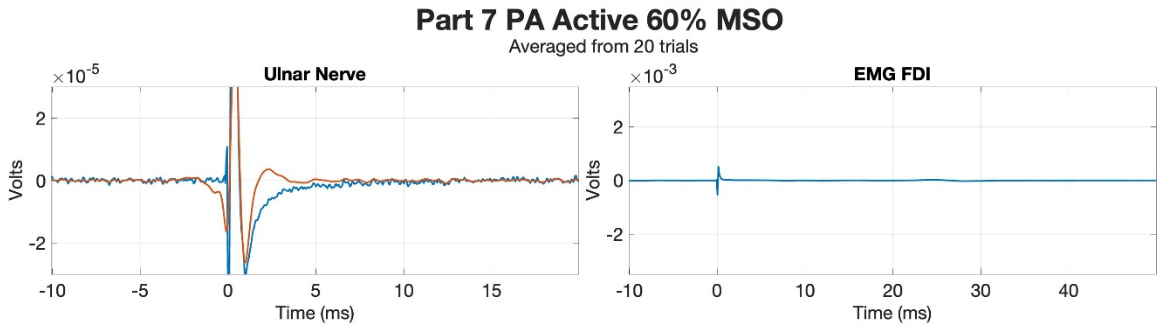
Part 7 PA Active 60% MSO
Averaged from 20 trials
Ulnar Nerve
EMG FDI
Volts
Time (ms)

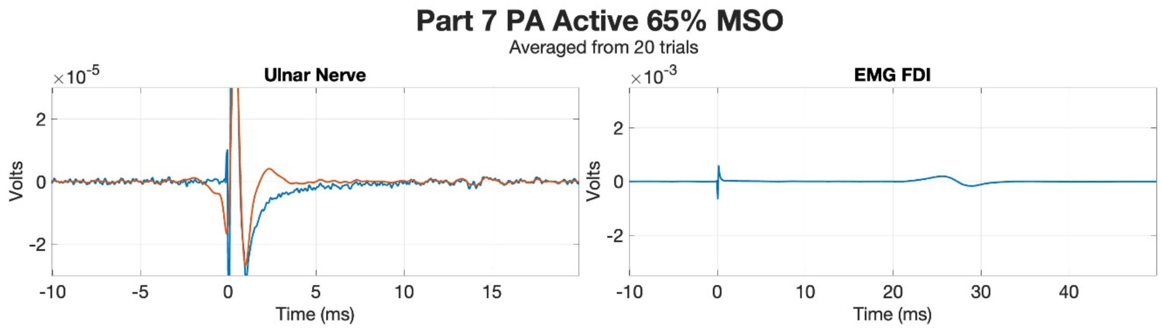
Part 7 PA Active 65% MSO
Averaged from 20 trials
Ulnar Nerve
EMG FDI
Volts
Time (ms)

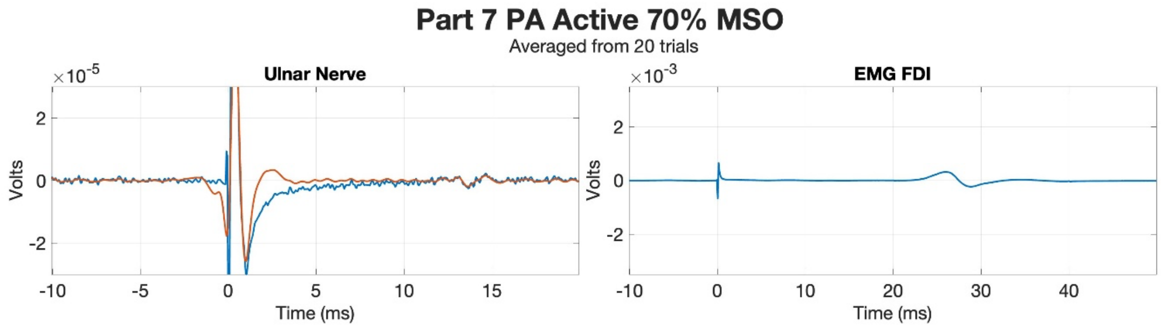
Part 7 PA Active 70% MSO
Averaged from 20 trials
Ulnar Nerve
EMG FDI
Volts
Time (ms)

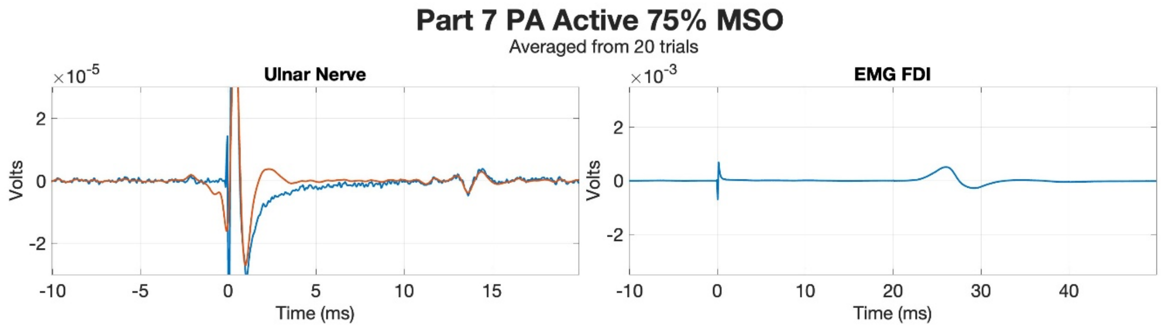
Part 7 PA Active 75% MSO
Averaged from 20 trials
Ulnar Nerve
EMG FDI
Volts
Time (ms)

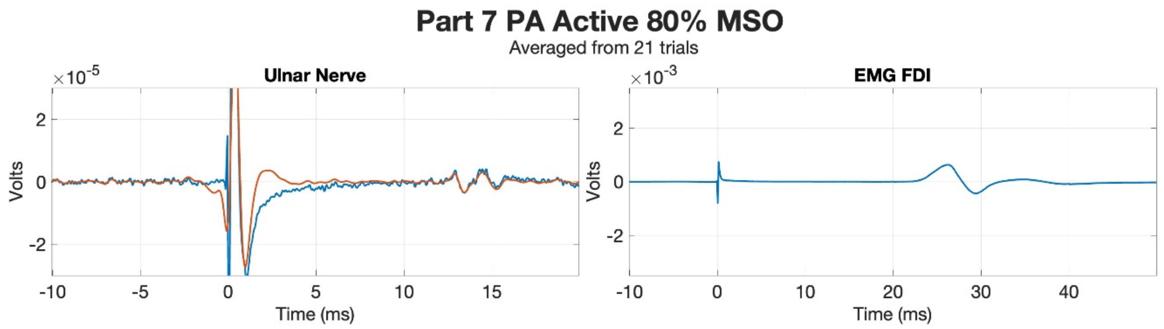
Part 7 PA Active 80% MSO
Averaged from 21 trials
Ulnar Nerve
EMG FDI
×10^-5
×10^-3
Volts
Time (ms)

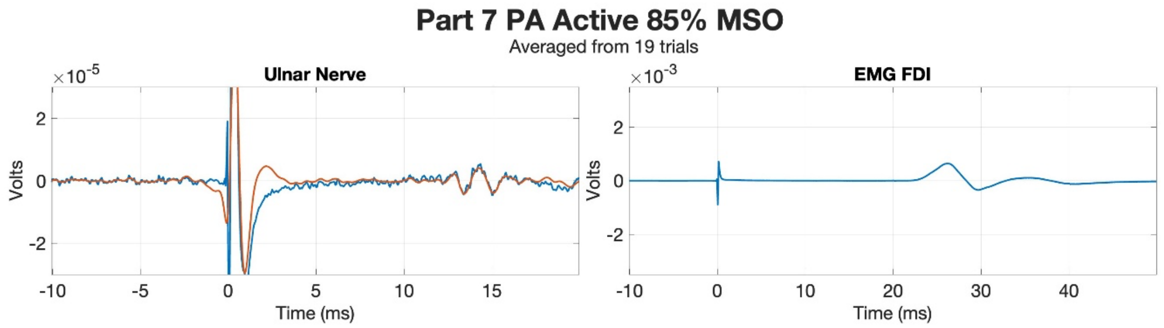
Part 7 PA Active 85% MSO
Averaged from 19 trials
Ulnar Nerve
EMG FDI
×10^-5
×10^-3
Volts
Time (ms)

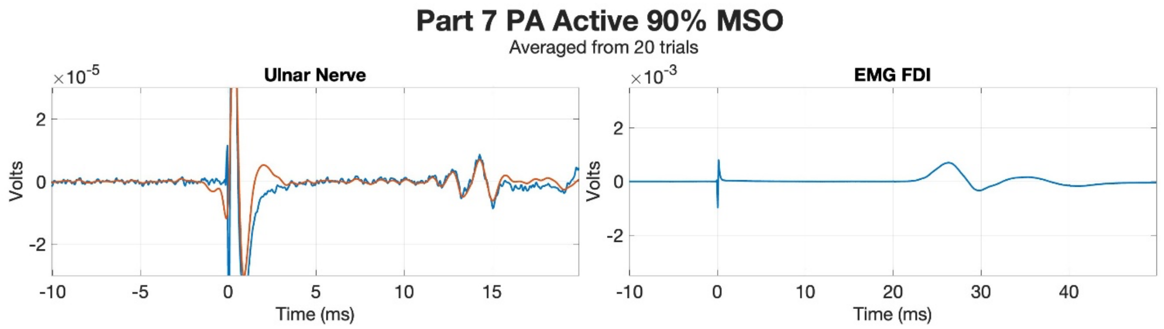
Part 7 PA Active 90% MSO
Averaged from 20 trials
Ulnar Nerve
EMG FDI
×10^-5
×10^-3
Volts
Time (ms)

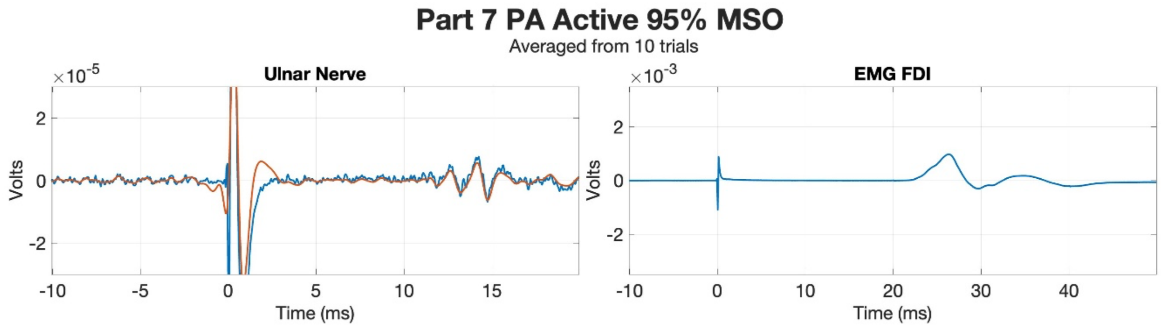
Part 7 PA Active 95% MSO
Averaged from 10 trials
Ulnar Nerve
EMG FDI
×10^-5
×10^-3
Volts
Time (ms)

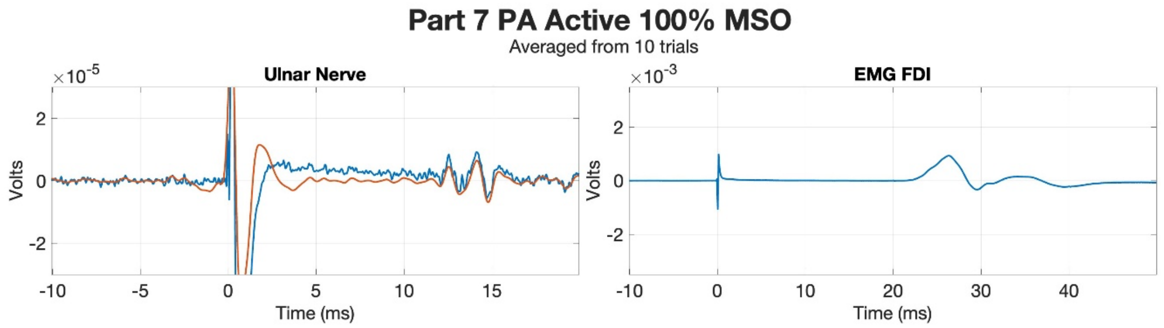
Part 7 PA Active 100% MSO
Averaged from 10 trials
Ulnar Nerve
EMG FDI
×10^-5
×10^-3
Volts
Time (ms)

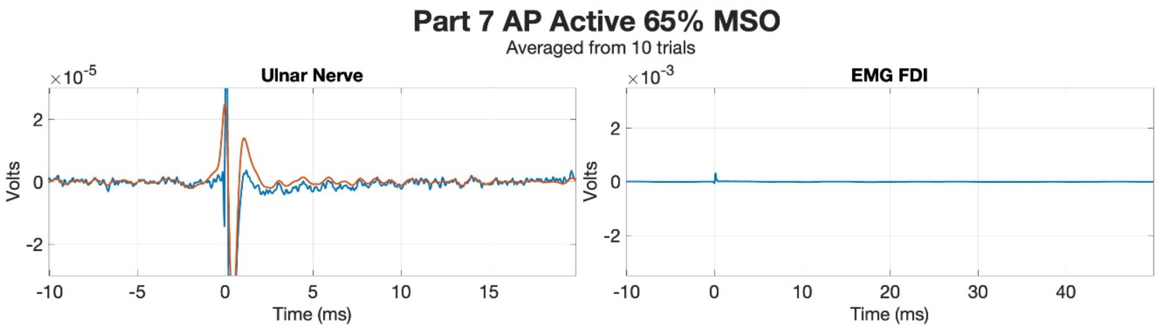
Part 7 AP Active 65% MSO
Averaged from 10 trials
Ulnar Nerve
EMG FDI
Volts
Time (ms)

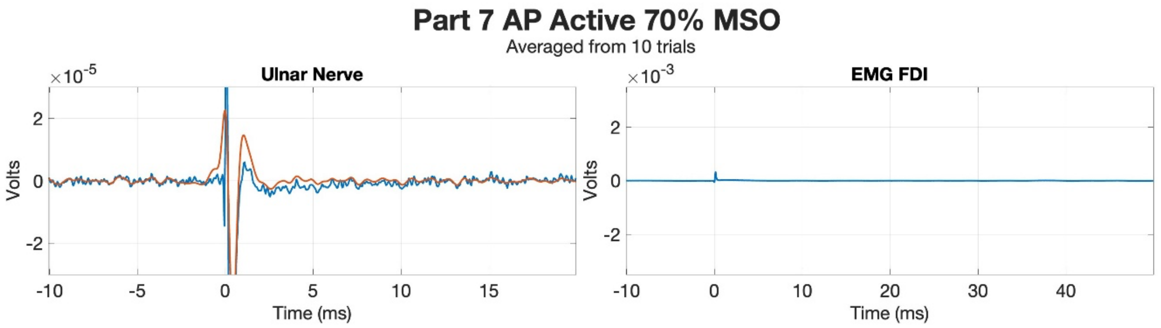
Part 7 AP Active 70% MSO
Averaged from 10 trials
Ulnar Nerve
EMG FDI
Volts
Time (ms)

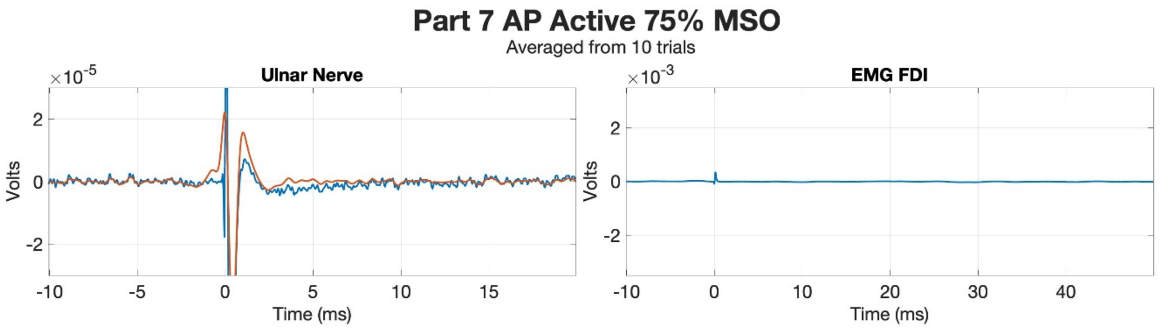
Part 7 AP Active 75% MSO
Averaged from 10 trials
Ulnar Nerve
EMG FDI
Volts
Time (ms)

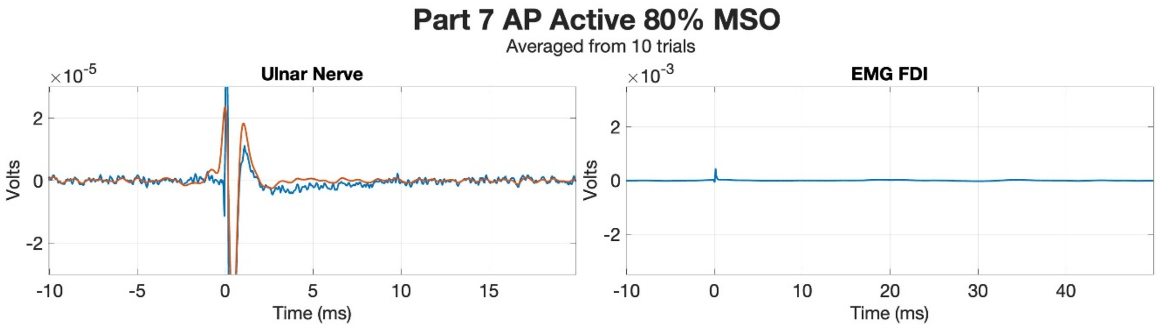
Part 7 AP Active 80% MSO
Averaged from 10 trials
Ulnar Nerve
EMG FDI
Volts
Time (ms)

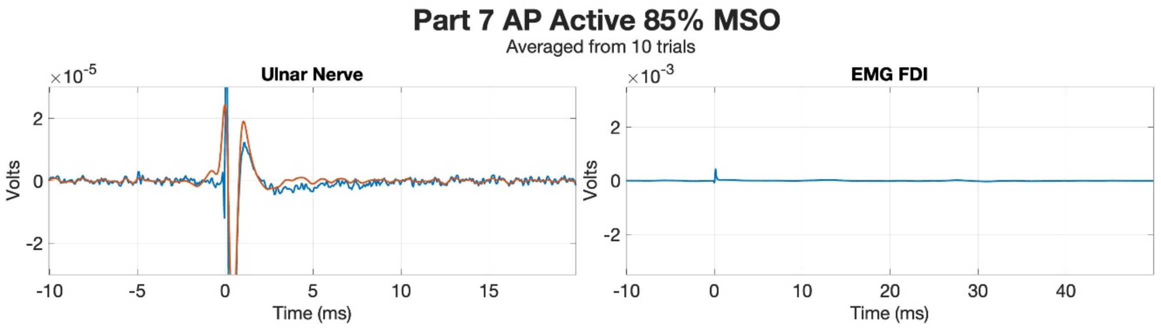
Part 7 AP Active 85% MSO
Averaged from 10 trials
Ulnar Nerve
EMG FDI
Volts
Time (ms)

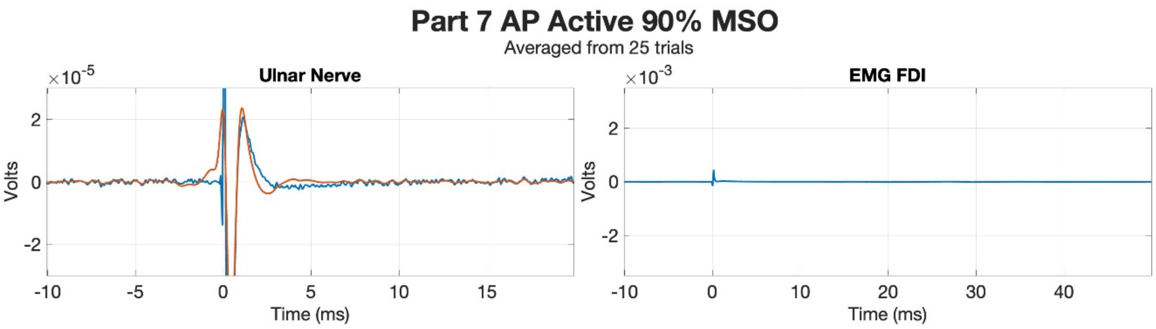
Part 7 AP Active 90% MSO
Averaged from 25 trials
Ulnar Nerve
EMG FDI
×10^-5
×10^-3
Volts
Time (ms)

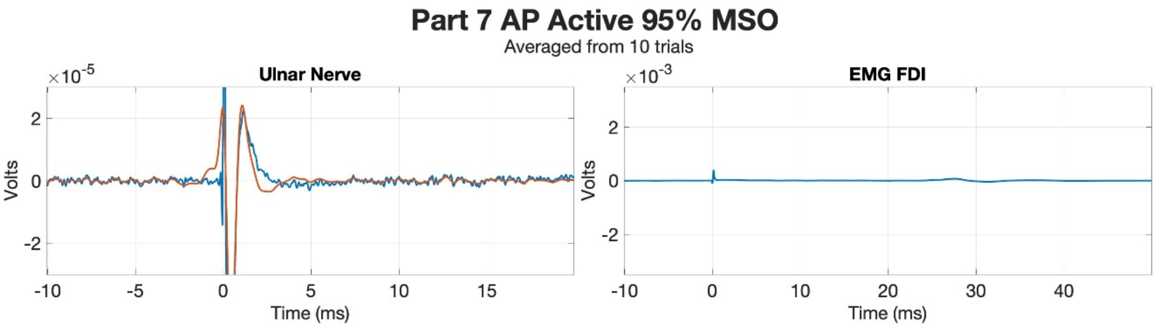
Part 7 AP Active 95% MSO
Averaged from 10 trials
Ulnar Nerve
EMG FDI
×10^-5
×10^-3
Volts
Time (ms)

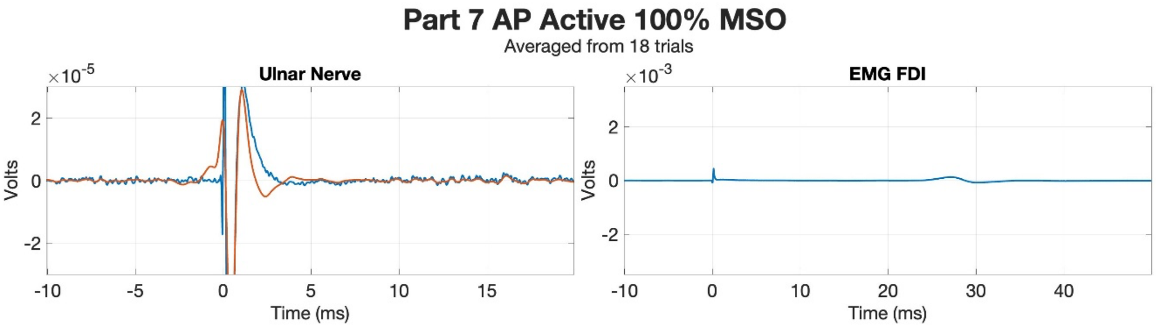
Part 7 AP Active 100% MSO
Averaged from 18 trials
Ulnar Nerve
EMG FDI
×10^-5
×10^-3
Volts
Time (ms)

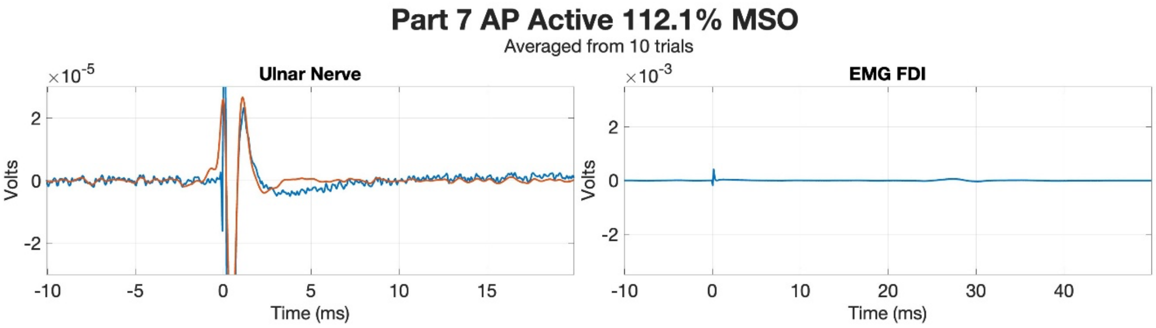
Part 7 AP Active 112.1% MSO
Averaged from 10 trials
Ulnar Nerve
EMG FDI
×10^-5
×10^-3
Volts
Time (ms)

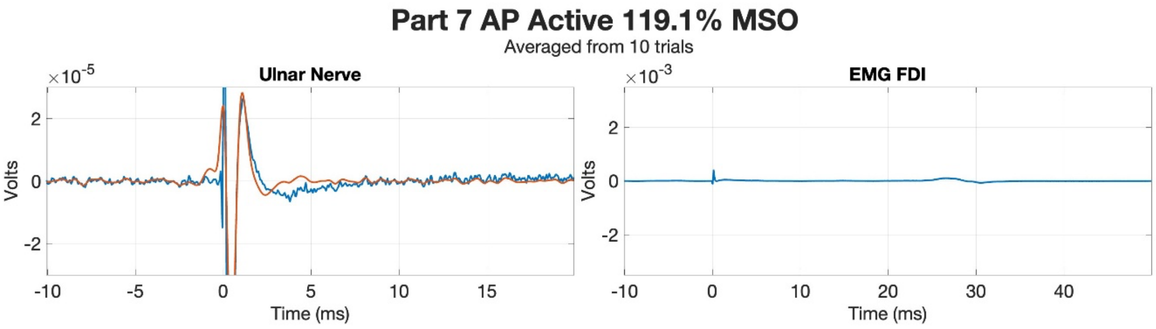
Part 7 AP Active 119.1% MSO
Averaged from 10 trials
Ulnar Nerve
EMG FDI
×10^-5
×10^-3
Volts
Time (ms)

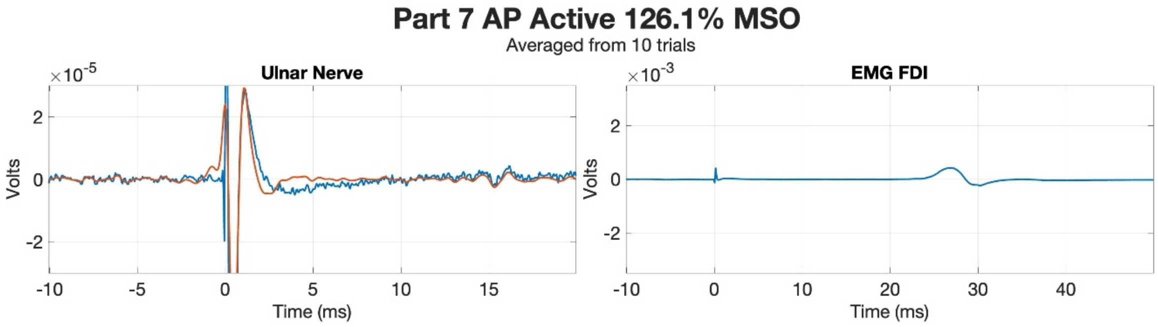
Part 7 AP Active 126.1% MSO
Averaged from 10 trials
Ulnar Nerve
EMG FDI
×10^-5
×10^-3
Volts
Time (ms)

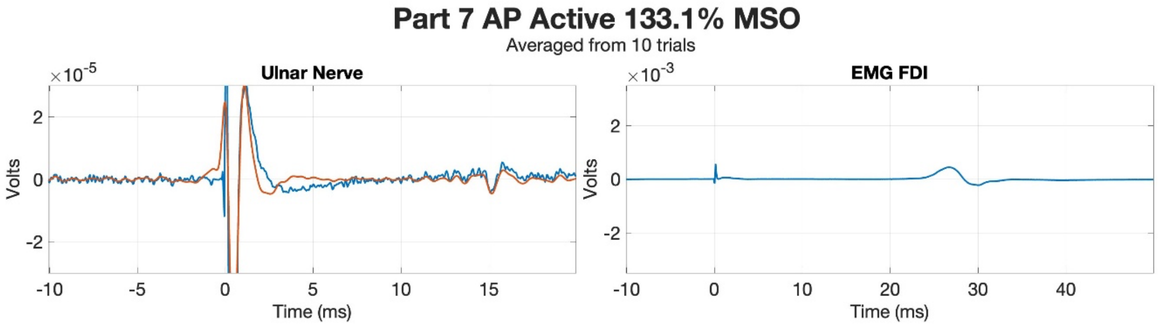
Part 7 AP Active 133.1% MSO
Averaged from 10 trials
Ulnar Nerve
EMG FDI
×10^-5
×10^-3
Volts
Time (ms)

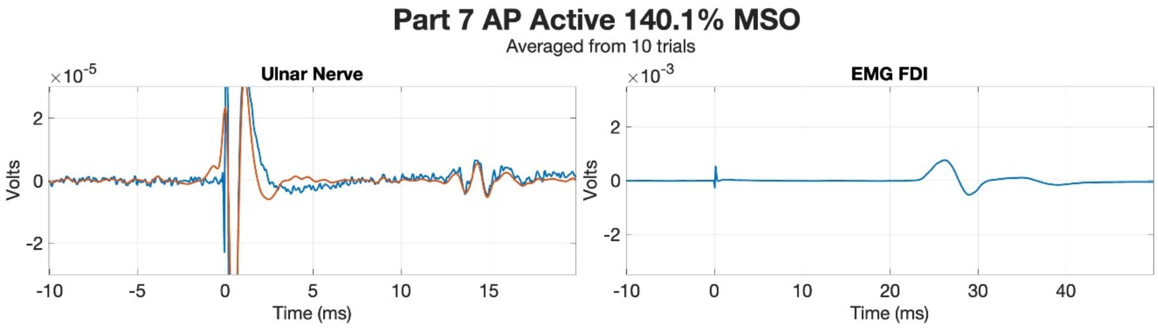
Part 7 AP Active 140.1% MSO
Averaged from 10 trials
Ulnar Nerve
EMG FDI
×10^-5
×10^-3
Volts
Time (ms)

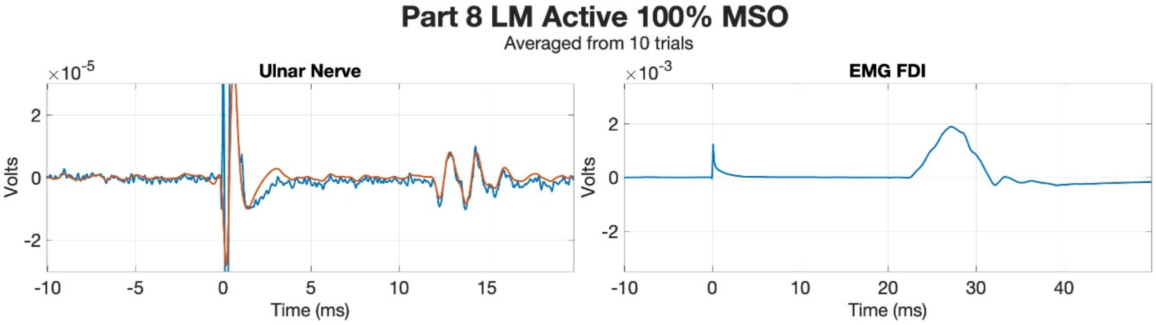
Part 8 LM Active 100% MSO
Averaged from 10 trials
Ulnar Nerve
EMG FDI
Volts
Time (ms)
×10^-5
×10^-3

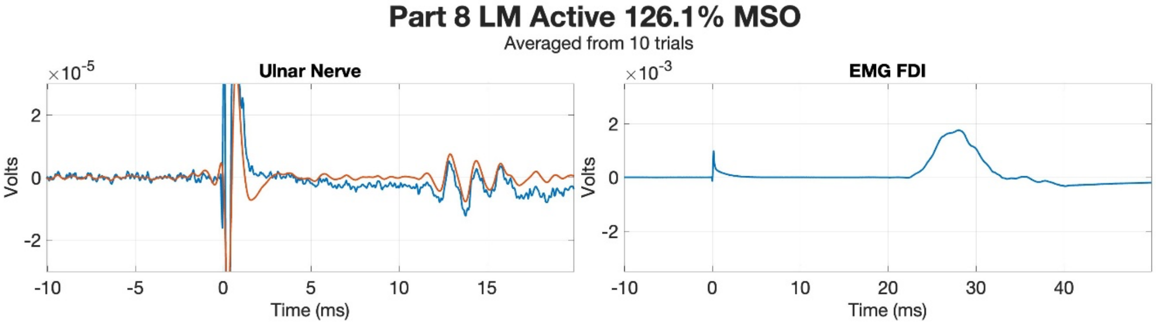
Part 8 LM Active 126.1% MSO
Averaged from 10 trials
Ulnar Nerve
EMG FDI
Volts
Time (ms)
×10^-5
×10^-3

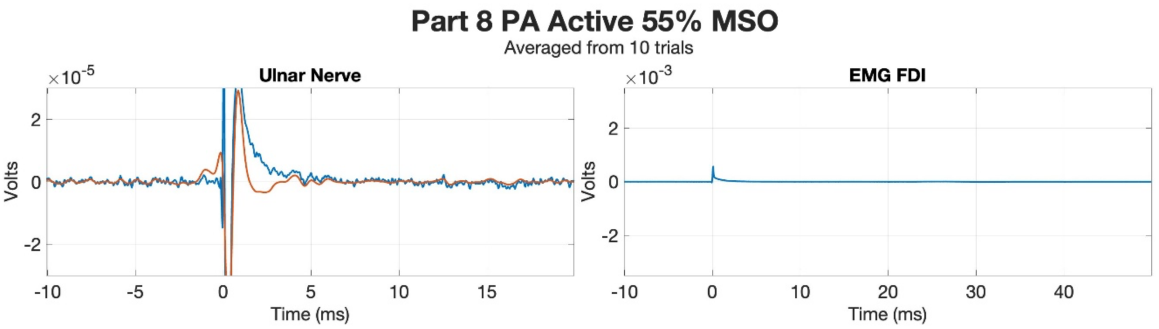
Part 8 PA Active 55% MSO
Averaged from 10 trials
Ulnar Nerve
EMG FDI
Volts
Time (ms)
×10^-5
×10^-3

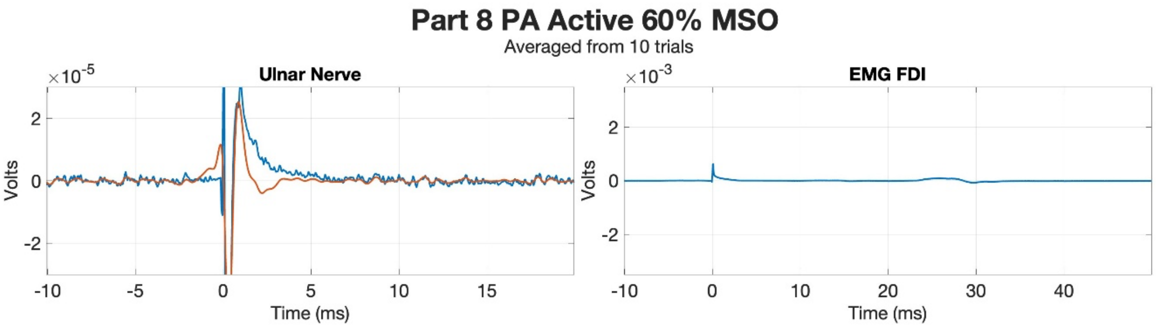
Part 8 PA Active 60% MSO
Averaged from 10 trials
Ulnar Nerve
EMG FDI
Volts
Time (ms)
×10^-5
×10^-3

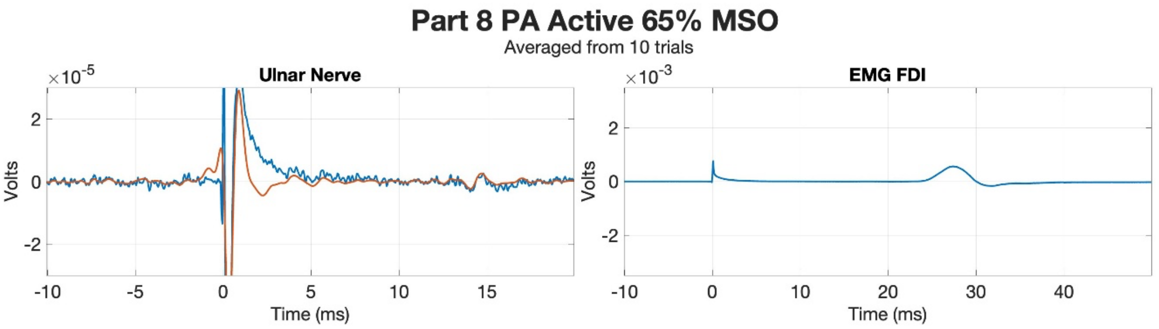
Part 8 PA Active 65% MSO
Averaged from 10 trials
Ulnar Nerve
EMG FDI
Volts
Time (ms)
×10^-5
×10^-3

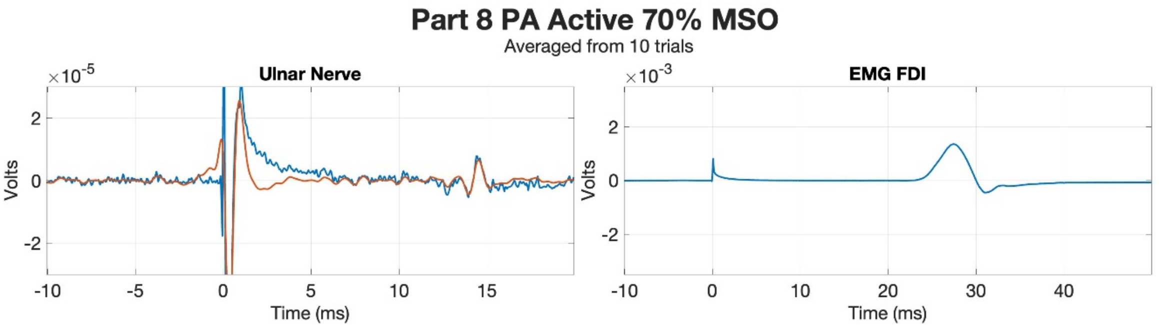
Part 8 PA Active 70% MSO
Averaged from 10 trials
Ulnar Nerve
EMG FDI
Volts
Time (ms)
×10^-5
×10^-3

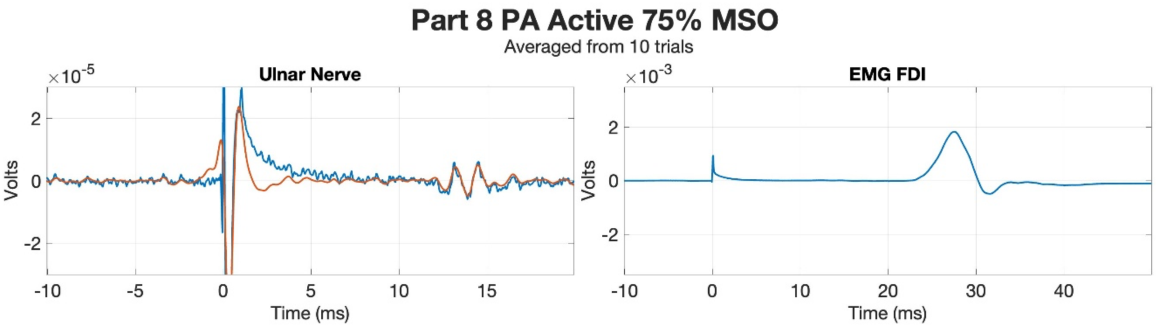
Part 8 PA Active 75% MSO
Averaged from 10 trials
Ulnar Nerve
EMG FDI
Volts
Time (ms)
×10^-5
×10^-3

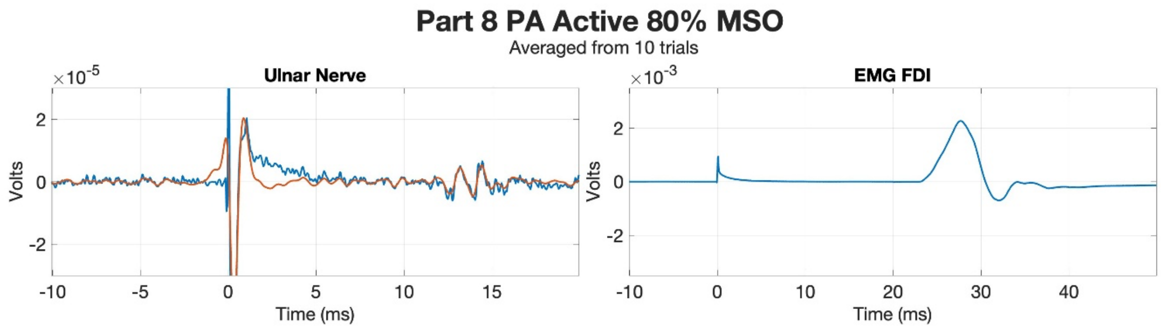
Part 8 PA Active 80% MSO
Averaged from 10 trials
Ulnar Nerve
EMG FDI
Volts
Time (ms)

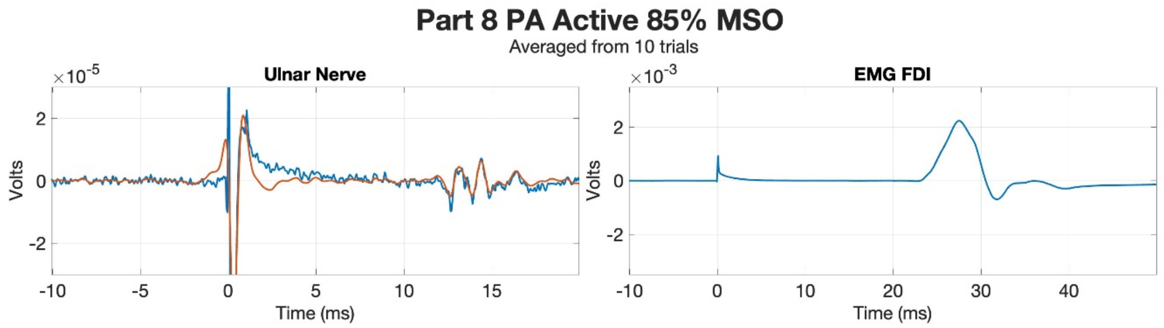
Part 8 PA Active 85% MSO
Averaged from 10 trials
Ulnar Nerve
EMG FDI
Volts
Time (ms)

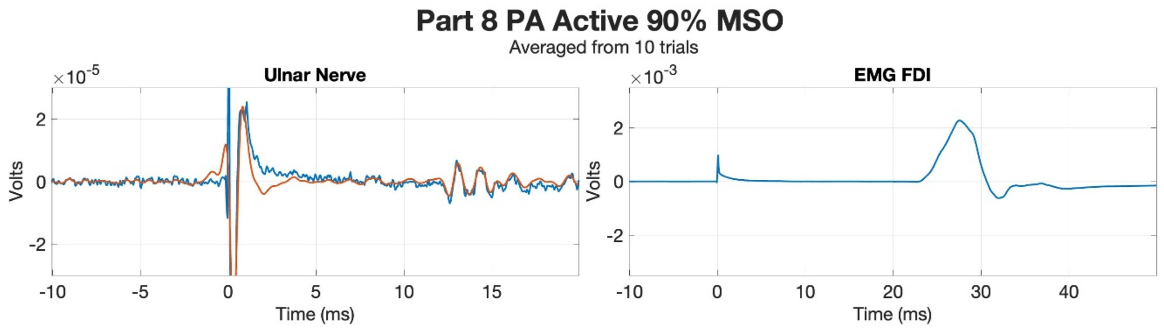
Part 8 PA Active 90% MSO
Averaged from 10 trials
Ulnar Nerve
EMG FDI
Volts
Time (ms)

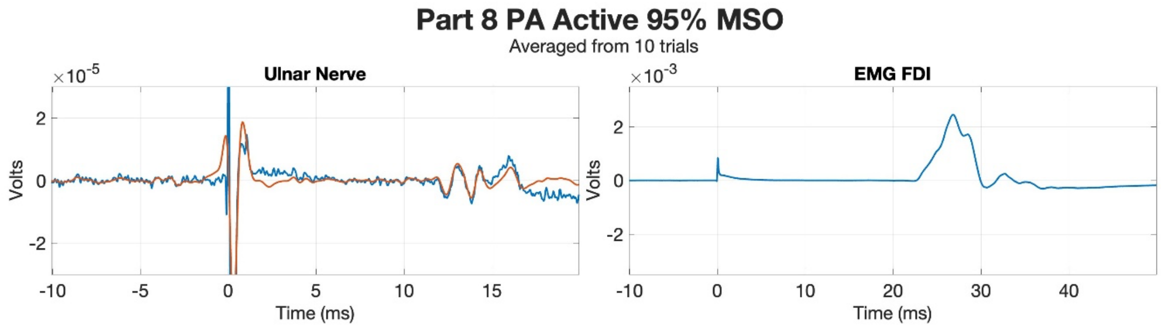
Part 8 PA Active 95% MSO
Averaged from 10 trials
Ulnar Nerve
EMG FDI
Volts
Time (ms)

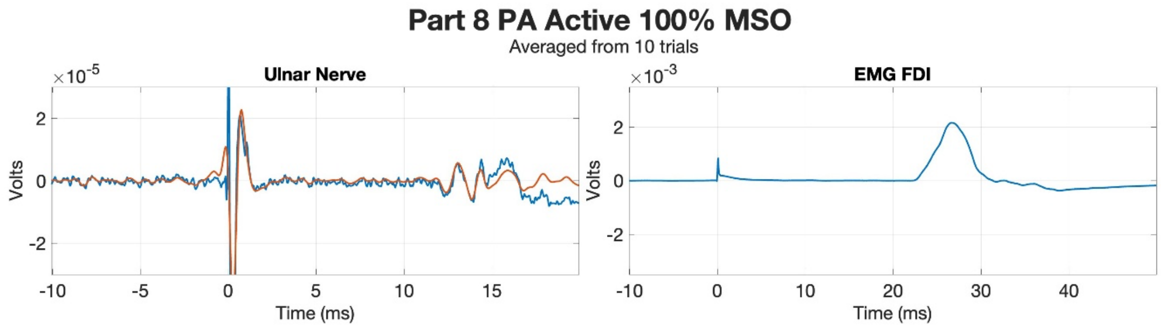
Part 8 PA Active 100% MSO
Averaged from 10 trials
Ulnar Nerve
EMG FDI
Volts
Time (ms)

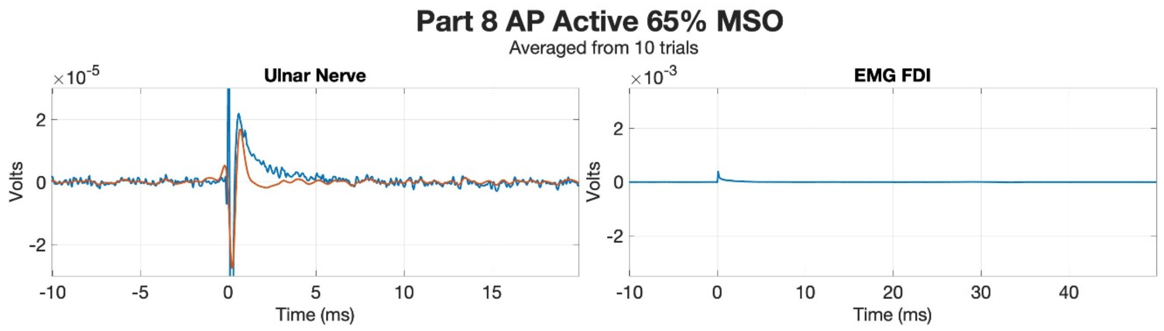
Part 8 AP Active 65% MSO
Averaged from 10 trials
Ulnar Nerve
EMG FDI
$\times 10^{-5}$
$\times 10^{-3}$
Volts
Time (ms)

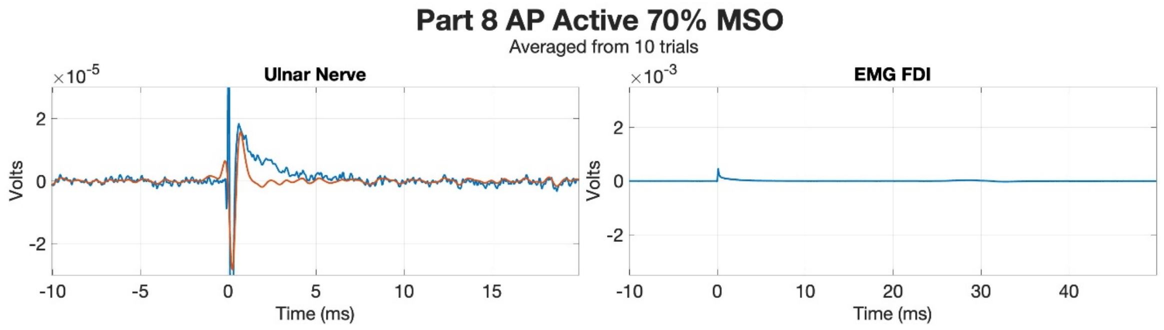
Part 8 AP Active 70% MSO
Averaged from 10 trials
Ulnar Nerve
EMG FDI
$\times 10^{-5}$
$\times 10^{-3}$
Volts
Time (ms)

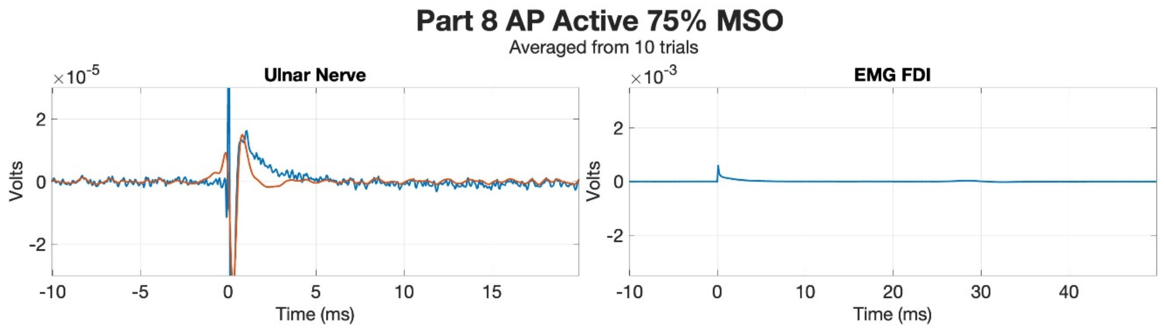
Part 8 AP Active 75% MSO
Averaged from 10 trials
Ulnar Nerve
EMG FDI
$\times 10^{-5}$
$\times 10^{-3}$
Volts
Time (ms)

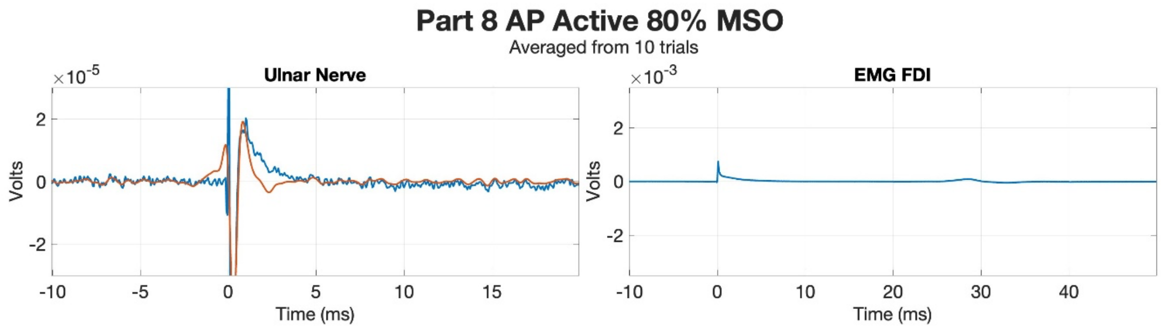
Part 8 AP Active 80% MSO
Averaged from 10 trials
Ulnar Nerve
EMG FDI
$\times 10^{-5}$
$\times 10^{-3}$
Volts
Time (ms)

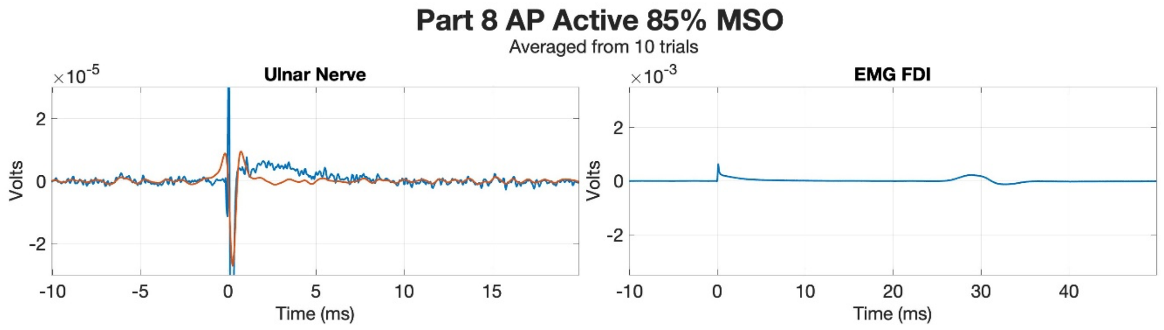
Part 8 AP Active 85% MSO
Averaged from 10 trials
Ulnar Nerve
EMG FDI
$\times 10^{-5}$
$\times 10^{-3}$
Volts
Time (ms)

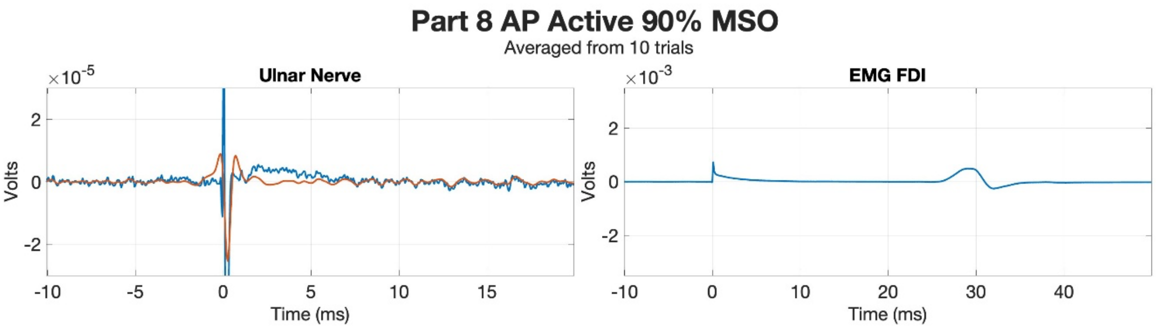
Part 8 AP Active 90% MSO
Averaged from 10 trials
Ulnar Nerve
×10^-5
Volts
Time (ms)
EMG FDI
×10^-3
Volts
Time (ms)

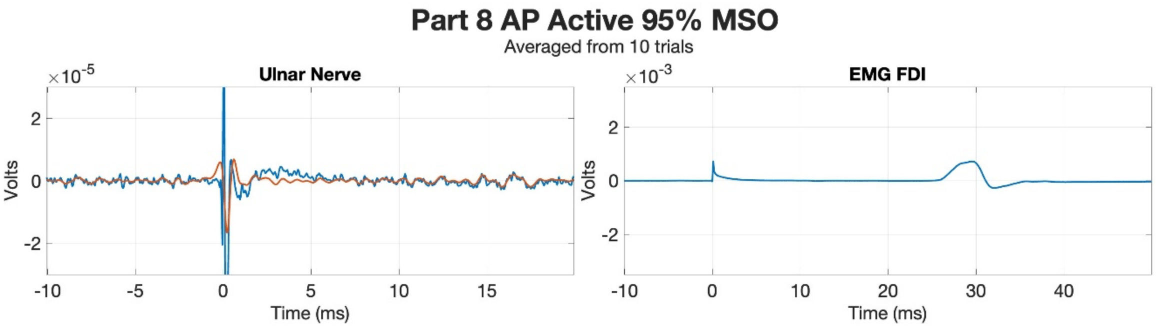
Part 8 AP Active 95% MSO
Averaged from 10 trials
Ulnar Nerve
×10^-5
Volts
Time (ms)
EMG FDI
×10^-3
Volts
Time (ms)

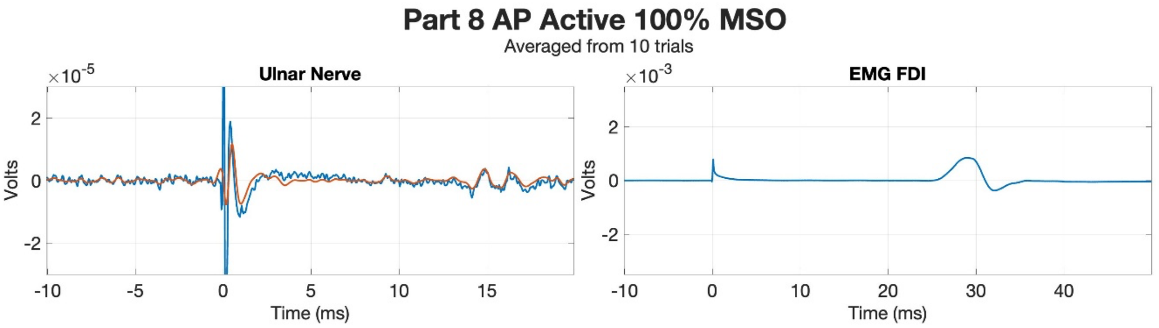
Part 8 AP Active 100% MSO
Averaged from 10 trials
Ulnar Nerve
×10^-5
Volts
Time (ms)
EMG FDI
×10^-3
Volts
Time (ms)
